# Peptidoglycan maturation controls spatiotemporal organisation of outer membrane proteins in *Escherichia coli*

**DOI:** 10.1101/2022.04.11.487844

**Authors:** Gideon Mamou, Federico Corona, Ruth Cohen-Khait, Dawei Sun, Pooja Sridhar, Timothy J. Knowles, Colin Kleanthous, Waldemar Vollmer

## Abstract

Linkages between the outer membrane of Gram-negative bacteria and the peptidoglycan layer are crucial to the maintenance of cellular integrity and enable survival in challenging environments^1–5^. The functionality of the outer membrane relies on outer membrane proteins (OMPs), which are inserted by the β-barrel assembly machine, BAM^6, 7^. Previous work has shown that growing *Escherichia coli* cells segregate old OMPs towards the poles by an unknown mechanism^8^. Here, we demonstrate that peptidoglycan underpins the spatiotemporal organisation of OMPs. Mature, tetrapeptide-rich peptidoglycan binds to BAM components and suppresses OMP foldase activity. Nascent peptidoglycan, which is enriched in pentapeptides and concentrated at septa^9^, associates with BAM poorly and has little impact on its activity, leading to preferential insertion of OMPs at division sites. Synchronising OMP biogenesis to cell wall growth enables bacteria to replenish their OMPs by binary partitioning. Our study reveals that Gram-negative bacteria coordinate the assembly of two major cell envelope layers by rendering OMP biogenesis responsive to peptidoglycan maturation. This coordination offers new possibilities for the design of antibiotics that disrupt cell envelope integrity.

## Main

Insertion of outer membrane β-barrel proteins in Gram-negative bacteria is catalysed by the BAM complex^6, 7^, comprising the β-barrel protein BamA and four accessory lipoproteins, BamBCDE^10–12^. BamA is essential for viability which, in conjunction with its surface exposure, makes it a promising target for Gram-negative-specific antibiotics^13–16^. Much is known of the mechanism by which BamA catalyses the folding of OMPs *in vitro*^17–20^. By contrast, little is known of BamA-mediated OMP biogenesis in the asymmetric OM of live bacteria. Some studies suggest new OMPs are inserted preferentially at mid-cell^8, 21, 22^ while others suggest OMPs are inserted throughout the membrane^23, 24^. These contrasting views are confounded by recent super-resolution microscopy studies in fixed cells showing BamA clustered throughout the OM^25^.

### BAM activity is linked to the *E. coli* cell cycle

To determine where OMPs emerge and if their insertion in the OM is modulated, we first ascertained the localization of BamA in live *E. coli* cells. We visualised BamA by epifluorescence microscopy and 3D-structured illumination microscopy (SIM) following labelling with a specific, high-affinity monoclonal Fab antibody (MAB2)^13^ that binds extracellular loop 6 in BamA with no impact on growth (Extended Data Fig. 1A-C). We found that BamA clusters into small (average diameter ∼150 nm), uniformly distributed islands (8-10/μm^2^) on the cell surface (Extended Data Fig. 1D-F, Video 1), in agreement with data on fixed cells^25^. We next investigated where newly synthesised OMPs appear on the cell surface in relation to this distribution of BamA. We focused on two TonB-dependent transporters (TBDTs), the siderophore transporter FepA and the vitamin B_12_ transporter BtuB. Both TBDTs were labelled with high-affinity, fluorescently-labelled colicins which exploit these OMPs as receptors^8, 26^. ColB, specific for FepA, was fused to mCherry or GFP, and ColE9, specific for BtuB, was labelled with AF488. Both OMPs were separately expressed in *E. coli* from plasmids and induced with arabinose (Extended Data Fig. 2A-B & 3A). Time-course studies with FepA indicated that the OMP appeared on the bacterial surface after ∼3 min (Extended Data Fig. 2C), and so all subsequent experiments included at least this time for induction. Within populations of cells that had been induced to express either FepA or BtuB, two clear zones of OMP biogenesis were observed; predominant were sites of cell division - yielding labelled septa and cells with unipolar labelling - and, to a lesser degree, OMPs on the long axis of cells (Fig. 1A-B, Video 2). Little or no OMP biogenesis was observed at old poles (Extended Data Fig. 2D-E, 3B-D), in agreement with previous reports^8, 21^. We next co-labelled BamA and either FepA or BtuB, the latter following a brief period of induction. Co-labelling revealed a clear divergence between BamA localisation and the sites of OMP insertion (Extended Data Fig. 3E). While BamA is distributed uniformly across the entire cell surface, including the poles, OMP biogenesis is confined to the long axis of cells and division sites (Fig. 1C-D). These data demonstrate conclusively that BamA is not equally active in the OM but that its OMP insertion activity is modulated in a cell cycle dependent manner.

**Figure 1.**
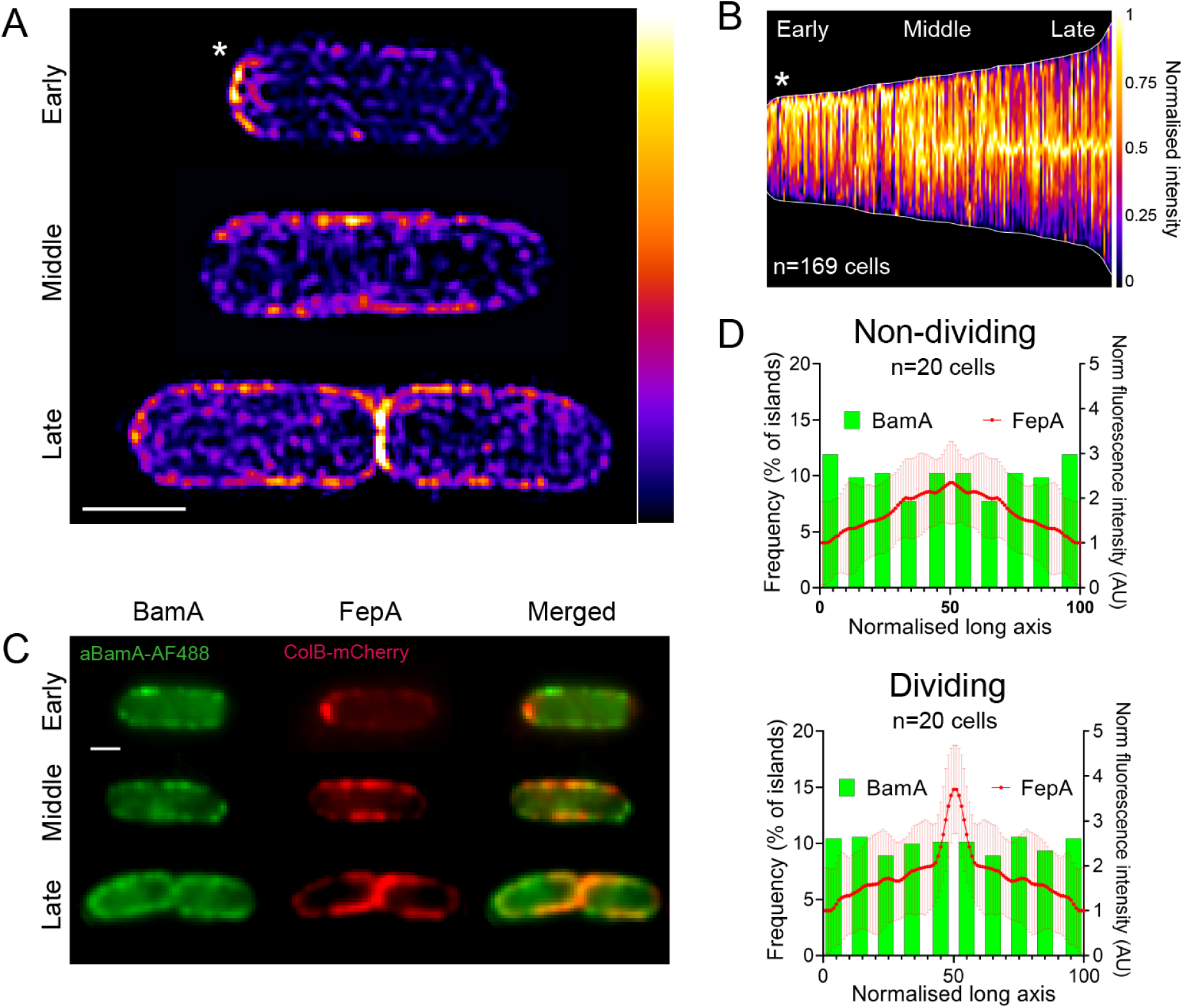
OMP biogenesis mirrors the cell cycle and not localisation of BamA in the OM of live *E. coli* cells. **(A)** Biogenesis patterns for FepA, stained with ColB-GFP, following a 5 min- induction (0.4% arabinose). Shown are heatmaps of individual cells representing different cell cycle stages. **(B)** Demograph showing normalised fluorescence intensity across multiple cells following 5 min induction of FepA biogenesis. Cells are aligned to show the more intense pole at the top. **(C)** Co-labelling of BamA and FepA at different cell cycle stages following 7 min of FepA induction. **(D)** Comparison of FepA biogenesis regions (7 min induction; ± SD, *red line*) with the distribution of BamA-containing islands (*green columns*) in dividing and non-dividing *E. coli* cells. Scale bar, 1 μm.

### OMP biogenesis co-localizes with cell wall synthesis

Peptidoglycan (PG) is made of glycan chains connected by short peptides and forms a thin, net-like layer, called saculus, underneath the outer membrane. The patterns of OMP biogenesis are reminiscent of those seen previously for PG in *E. coli*^27, 28^. We therefore investigated the correlation between PG and OMP biogenesis through co-labelling experiments. PG biogenesis was imaged following incorporation of the fluorescent D-amino acid HADA^29^ (Extended Data Fig. 4A-C, Video 3) and OMP biogenesis by labelling induced FepA with ColB-GFP. Cells were incubated for 7 min with both HADA and arabinose for *fepA* expression, fixed and surface-exposed FepA labelled with ColB-GFP. Similar patterns of fluorescence labelling were observed, with particularly strong labelling for both PG and OMP at division sites (Fig. 2A-C). Co-localisation analysis of fluorophore pixel intensity across cells also revealed a strong correlation between PG and OMP fluorescence and this correlation was significantly greater than that for BamA and OMP fluorescence (Extended Data Fig. 4D). Closer inspection of labelling at division septa revealed they segregated into two populations; those with both PG and OMP labelling (group 2) and those with only PG label (group 1) (Fig 2B). No cells were observed that had only OMP labelling at division sites. Fluorescence scans across each cell type and demographs of cell populations (Fig. 2B, C) demonstrated that PG biogenesis always appeared more advanced than OMP biogenesis. In addition, septal widths were narrower for group 2 cells compared to group 1 cells (Fig 2B). While the interpretation of these differences is complicated by the distinct, multi-step biosynthetic routes that result in label incorporation, they suggest that cell wall biogenesis precedes the emergence of OMPs at division sites. Dual labelling experiments that included an additional (3 min) pre-induction period for *fepA* expression prior to the addition of HADA generated patterns of PG/OMP labelling similar to those without pre-induction (Extended Data Fig. 4E-G). We conclude that the patterns of OMP and PG biogenesis closely mirror each other in exponentially growing cells suggesting extensive cross-talk between these layers of the bacterial cell envelope.

**Figure 2.**
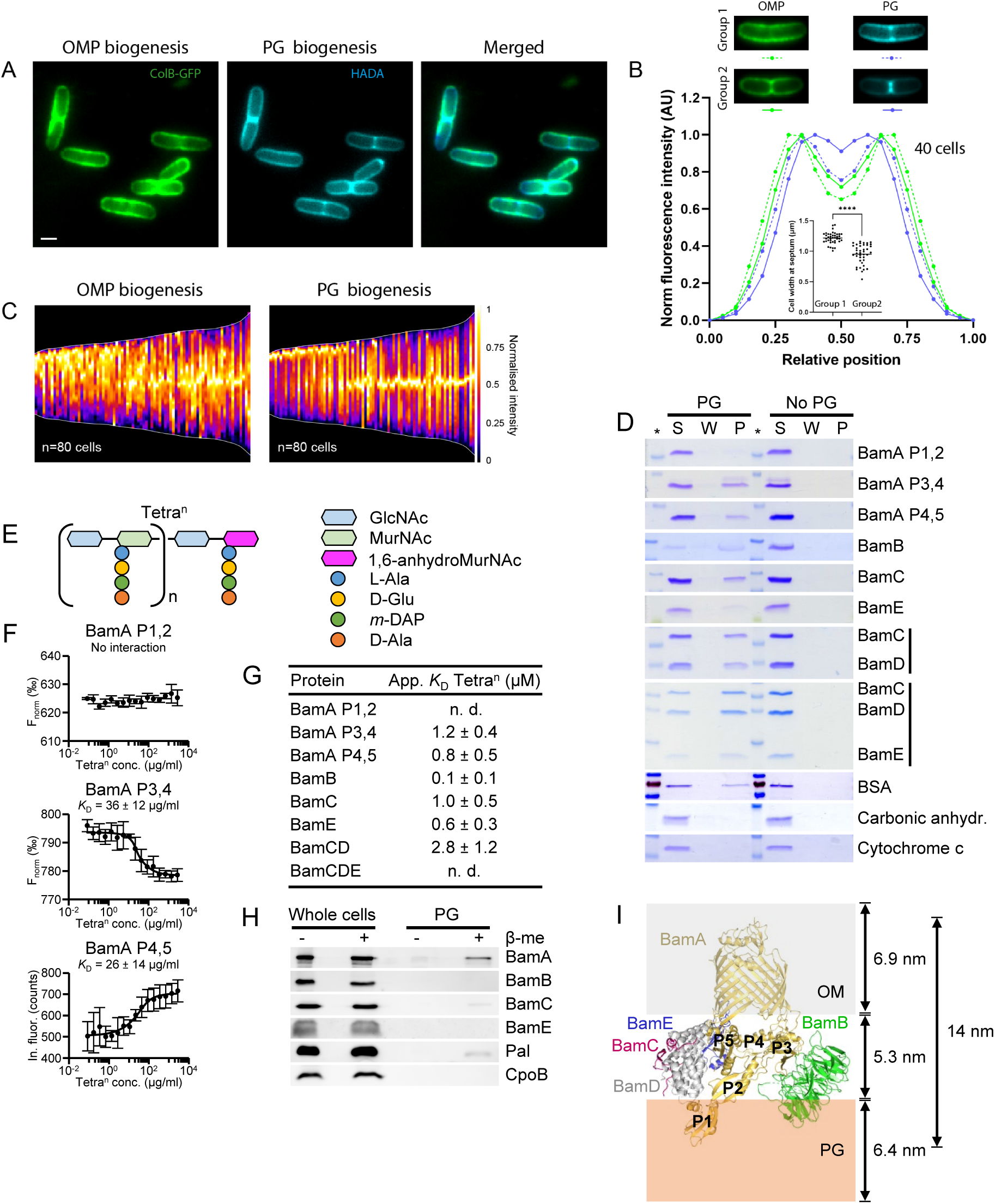
The cell wall plays a pivotal role in OMP biogenesis. **(A)** Co-labelling of PG (HADA) and the OMP FepA (ColB-GFP). PG staining and FepA induction (0.4% arabinose) were carried out simultaneously for 7 min. **(B)** Comparison of the fluorescence intensity profiles of PG (HADA) and OMP biogenesis (FepA, ColB-GFP) across the septum of dividing cells. Shown are profiles of cells at two different stages of septum formation (group1/group2). Inset displays the width of individual cells at the designated division plane. **(C)** Demographs comparing the normalised fluorescence distribution of FepA and PG biogenesis in multiple cells. HADA and FepA induction were carried out as in A. **(D)** Interaction of BAM proteins with isolated PG *in vitro* analysed by SDS-PAGE and Coomassie Blue staining. *S*, supernatant fraction; *W*, wash fraction; *P*, pellet fraction. **(E)** Schematic representation of poly-disaccharide-tetrapeptide chains (Tetra^n^). **(F)** Interaction of POTRA domains of BamA with Tetra^n^ measured by MST. **(G)** Apparent *K*_D_ values (µM) for the interaction of Bam proteins or Bam sub-complexes with Tetra^n^ measured by MST. Values are mean ± SD of three independent experiments (F and G). **(H)** Interaction of BamA and BamC with PG in *E. coli* MC1061 cells, detected by specific antibodies after *in vivo* cross-linking. Pal and CpoB were used as positive and negative controls, respectively. β-mercaptoethanol (β-me) reverses the cross-links releasing proteins from PG. **(I)** Schematic depiction of the position of the BAM complex (PDB code: 5AYW,^10^) relative to PG in the cell envelope. The approximate width and position of PG (orange, transparent), OM (grey, transparent) and OM-PG distance were modelled on measurements from Matias *et al*., 2003^47^.

### BAM proteins bind to PG *in vitro* and *in vivo*

We probed the link between OMP and PG biogenesis by first determining if purified BAM proteins physically interact with PG isolated from wild-type *E. coli* MC1061 in PG pull-down experiments^30, 31^. We tested three fragments from BamA encompassing POTRA domains 1-2 (BamA P1,2), 3-4 (BamA P3,4) and 4-5 (BamA P4,5), soluble versions of BamB, BamC and BamE, and BamCD and BamCDE complexes. We detected specific binding of all BAM proteins, except BamA P1,2, and the two sub-complexes to PG (Fig. 2D). We next tested the interaction of BAM proteins with soluble, uncross-linked disaccharide-tetrapeptide chains (Tetra^n^, Fig. 2E) by microscale thermophoresis (MST). We generated Tetra^n^ by digesting PG from *E. coli* BW25113Δ6LDT, which exclusively contains tetrapeptides and tetra-tetra (4-3) cross-links^29^, with the DD-endopeptidase MepM^32^ (Extended Data Fig. 5A). BamA P3,4 and BamA P4,5 but not BamA P1,2 interacted with Tetra^n^ (Fig. 2F). We also detected interaction of BamB, BamC, BamE and BamCD, but not BamCDE, with Tetra^n^, with apparent *K*_D_ values between 0.15 and 2.8 µM (Fig. 2G, Extended Data Fig. 5). There were no changes in the MST responses in the absence of Tetra^n^ (i.e., serial dilutions of mock MepM digestion without PG) (Extended Data Fig. 5). We conclude that BamA, BamB, BamC, BamE and BamCD bind specifically to Tetra^n^ chains of PG. To test whether BAM proteins interact with PG in *E. coli* cells, we treated *E. coli* MC1061 cultures with the chemical cross-linker DTSSP followed by isolation of PG, reversal of cross-linking and detection of Bam proteins by SDS-PAGE with specific antibodies^33^. We found BamA and BamC to be cross-linked to PG isolated from DTSSP-treated cells (Fig. 2H). BamB and BamE were not cross-linked to PG. As expected, the method identified the known PG-binding OM lipoprotein Pal^34^ whilst the lipoprotein CpoB, which does not interact with PG^35^, was not detected. These results indicate that BamA and BamC interact with PG in *E. coli* cells. BAM proteins are likely to be close to the cell wall based on dimensions of the complex from structural studies (Fig. 2I). The relative positions between them is likely to vary however due to the highly dynamic nature of the OMP folding cycle^36^, which may explain why binding of PG to some BAM components is detected *in vitro* but not *in vivo*.

### PG differentially impacts BAM activity *in vitro*

Sites of PG biosynthesis are characterised by a transient enrichment in pentapeptides, which are present in PG precursors but not in mature PG due to the action of DD-carboxypeptidases^9^ We therefore tested whether differences in PG composition might account for the cell-cycle dependence of OMP biogenesis. Specifically, we tested if differences in stem peptide composition between mature (tetrapeptide-rich) and nascent (pentapeptide-rich) PG affected the BAM-PG interaction. We purified the complete BAM complex (BamABCDE) and tested its binding to different PG sacculi preparations. We used MC1061 as a source for mature PG and the multiple DD-carboxypeptidase mutant CS703-1^37, 38^ as a source for pentapeptide-rich PG, mimicking nascent PG (Fig. 3A; Extended Data Fig. 6A). The BAM complex pulled down with tetrapeptide-rich PG, while we observed almost no binding to pentapeptide-rich PG (Fig. 3B). Purified BAM subunits also showed a reduced interaction with pentapeptide-rich PG in comparison with tetrapeptide-rich PG (Extended Data Fig. 6B). We conclude that BAM proteins have greater affinity for tetrapeptide-rich PG than pentapeptide-rich PG.

**Figure. 3.**
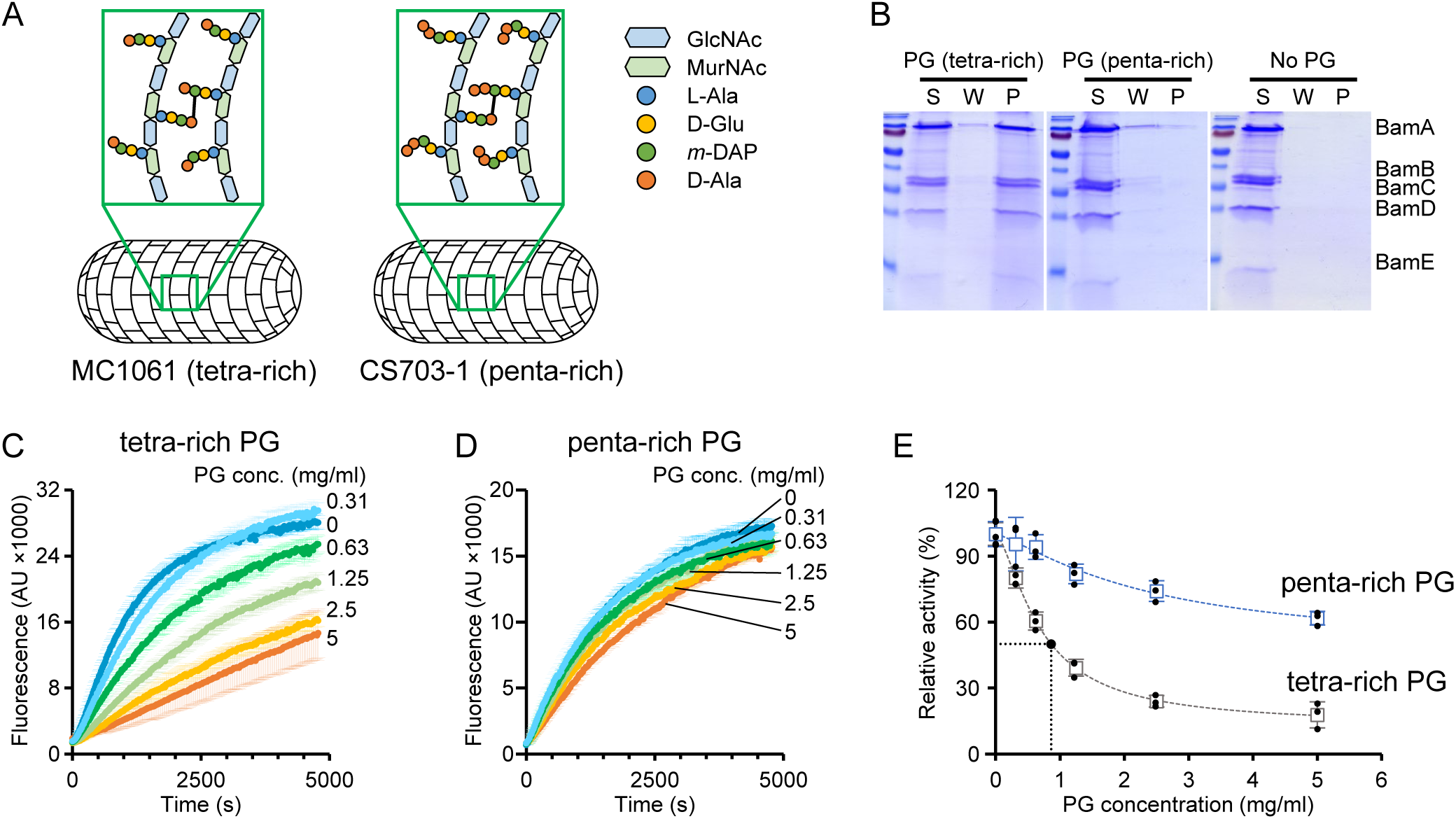
Nascent (pentapeptide-rich) and mature (tetrapeptide-rich) PG differentially impact BAM activity. **(A)** Schematic representation of PG structure in MC1061 and CS703-1, which are enriched in tetrapeptides and pentapeptides, respectively. **(B)** Interaction of BamABCDE with tetrapeptide-rich and pentapeptide-rich sacculi *in vitro* analysed after pull-down by SDS-PAGE and Coomassie Blue staining. No PG, negative control without PG. *S*, supernatant fraction; *W*, wash fraction; *P*, pellet fraction. **(C-E)** Differences in dose-dependent effect of tetrapeptide-rich and pentapeptide-rich PG on BAM-mediated OmpT assembly *in vitro*. Values are mean ± SD of three independent experiments.

We next asked whether the composition of the peptides in PG impacted BAM activity *in vitro* using an established assay that monitors the liposome incorporation of the β-barrel protease OmpT. The protease activity of folded OmpT is detected as fluorescence following cleavage of a self-quenched peptide substrate^39–41^ (Extended Data Fig. 6C). Addition of tetrapeptide-rich PG from MC1061 reduced BAM activity, while pentapeptide-rich PG from CS703-1 had little effect, which was consistent with the stronger binding of BAM proteins and the whole complex to PG (Fig. 3C-E). Control experiments indicated that the effects of PG on OMP folding activity were specific to BAM (Extended Data Fig. 6D-F). PG impact on BAM activity was dose-dependent, with an EC_50_ of 0.86 mg/ml for tetrapeptide-rich PG and >5 mg/ml for pentapeptide-rich PG (Fig. 3C-E). Finally, we demonstrated that Tetra^n^ had little effect on BAM activity (Extended Data Fig. 6G), suggesting cross-linked high-molecular weight forms of PG modulate BAM activity. Taken together, these results indicate that binding of BAM components to tetrapeptide-rich PG reduces OMP folding activity while pentapeptide-rich PG, which binds BAM poorly, has a correspondingly milder effect.

### PG modulates OMP patterning *in vivo*

Since mature and nascent PG have differential effects on the OMP folding activity of BAM *in vitro*, we hypothesized that cell wall composition is the driver of OMP patterning in the OM of *E. coli in vivo*. To test this hypothesis, we induced OMP (*fepA*) expression in a pentapeptide-rich mutant strain (CS703-1) and its tetrapeptide-rich parent (CS109). In both strains, FepA appeared on the cell surface shortly after arabinose induction (Extended Data Fig. 7A). However, while the distribution of FepA in the tetrapeptide-rich strain was similar to that observed in a wild-type (BW25113) strain, including mid-cell bias, the pentapeptide-rich strain displayed a defective biogenesis pattern with reduced mid-cell bias (Extended Data Fig. 7B-D). Furthermore, we found that OMP and PG biogenesis no longer mirrored one another as co-localisation was significantly reduced (Fig. 4A, B). Given the aberrant nature of OMP insertion in this and other PG mutant strains (see below), we developed a new live cell assay by which we could determine the effectiveness of mid-cell OMP insertion even in highly disrupted cell envelopes. This assay exploited the fact that several OMP genes in *E. coli*, including *fepA*, are highly expressed in late-exponential/stationary phase but suppressed in lag-phase/early exponential cells^42, 43^. We used the migration of FepA, expressed from its endogenous promoter, as cells revived from stationary phase as a proxy for mid-cell OMP insertion. (Fig. 4C). We used this assay to assess the impact of PG mutant strains on their ability to drive polar migration of FepA after first having ascertained that all strains expressed comparable levels of BamA (Extended Data Fig. 7E-F, 8B). We also determined that they displayed uniform (or near uniform) FepA in stationary phase (Extended Data Fig. 8A). Comparing FepA localisation in wild-type and the pentapeptide-rich mutant following revival demonstrated that FepA distribution was bipolar in the former but nearly uniform in the latter, indicative of defective OMP insertion at mid-cell in the cell wall mutant (Fig. 4D-E). Decreased polar migration was also observed when only *dacA* (encoding PBP5) was deleted (Fig. 4D, E). This strain (CS12-7^38^) contains higher than normal levels of pentapeptide PG, but the increase is more moderate than in CS703-1^44^. Importantly, restoration of FepA polar migration was achieved upon plasmid expression of PBP5 in the CS703-1 background (Fig. 4D, E, Extended Data Fig. 8B-D). This effect was retained in a derivative of CS703-1 lacking *lpp* (CS703-1Δ*lpp*, Extended Data Fig. 8E, 9A-B), indicating that it is not the lack of OM tethering to the PG which results in changes to OMP biogenesis patterns. Finally, we demonstrated that PBP5 expression in the CS703-1 background was sufficient to restore significant *E. coli* OM stability when cells were challenged with the detergent SDS (Fig 4F), demonstrating that the OM instability associated with PG mutants is rescued when the cell wall defect is corrected. Cumulatively, our data show that it is the local density of pentapeptides that underpins the biogenesis patterns of OMPs and, consequently, the stability of the outer membrane in *E. coli* since nascent pentapeptides do not dampen BAM OMP insertion activity as do matured tetrapeptides (Fig. 4G). Pentapeptides are absent in the PG of old poles but abundant at division sites whereas inhibitory tetrapeptides show the inverse distribution, explaining why little or no OMP biogenesis is seen ifn old polar regions of *E. coli*.

**Figure 4.**
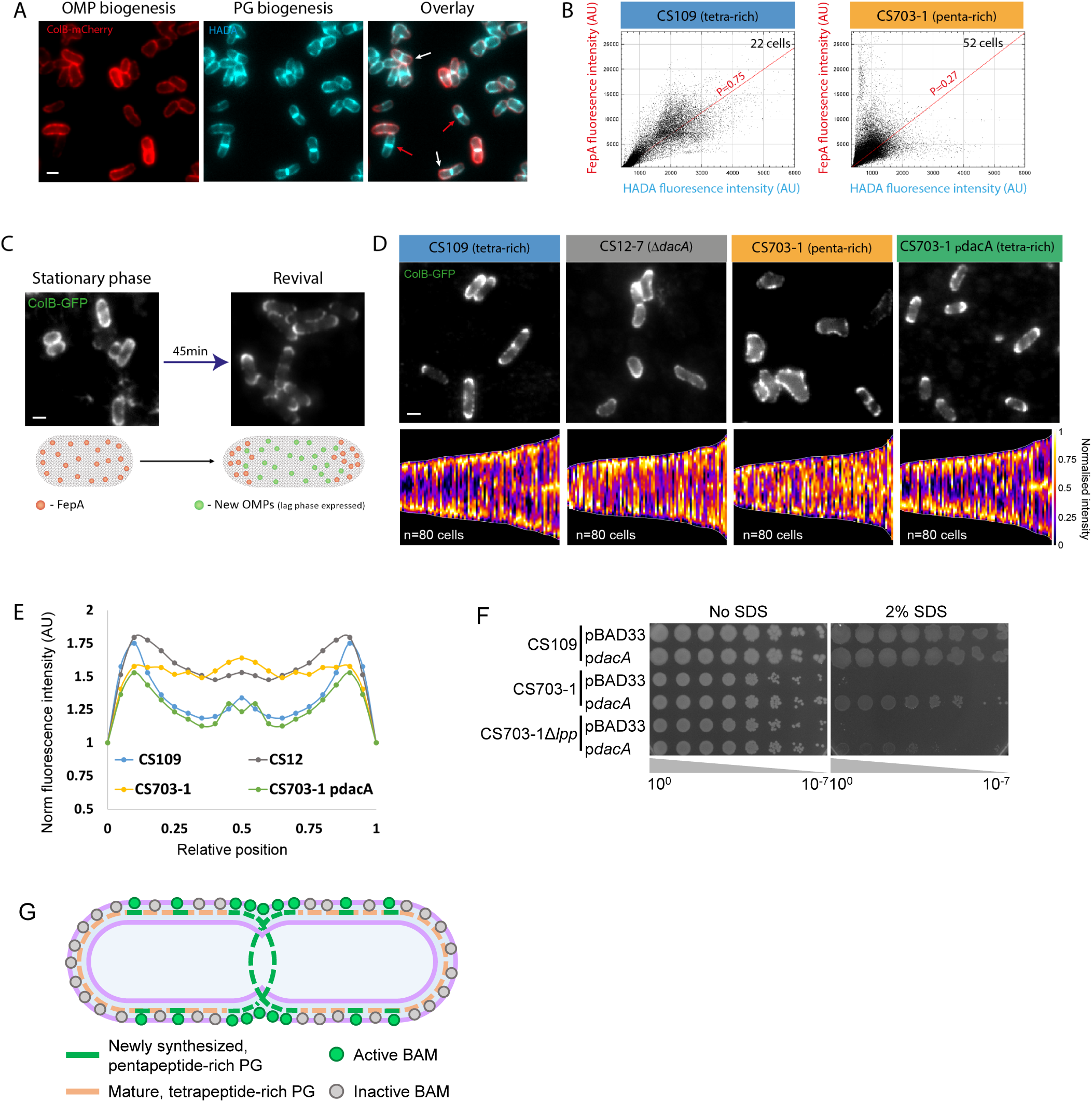
Cell wall composition affects the integrity and organization of the OM. **(A)** Co-labelling of PG and the OMP FepA in a pentapeptide rich strain (CS703-1). PG labelling (HADA) and FepA induction (0.4% arabinose) were initiated simultaneously for a total period of 7 min. Arrows indicate cells showing disparity between OMP and PG biogenesis **(B)** The fluorescence intensity of FepA vs HADA in tetra-rich (*right*) vs pentapeptide-rich (*left*) strains following co-labelling as in A. The representative pixel by pixel cytofluorograms of single images show that the strong correlation of OMP and PG biogenesis is abrogated in the pentapeptide-rich strain. **(C)** Polar migration assay whereby mid-cell OMP biogenesis bias can be discerned from the movement of stationary phase FepA as cells revive. Shown are representative images before/after resuspension in fresh media (*top*) and an illustration describing OMP movement during this period (*bottom*). **(D)** Polar migration of stationary phase FepA during revival. Shown are representative images (*top*) and demographs of normalised fluorescence intensities across multiple cells (*bottom*) 45 min after resuspension in fresh M9 media. The images and demographs show differently OMPs segregate between tetra-rich and penta-rich strains. **(E)** Distribution of FepA localisation along normalised cell lengths in the strains from D. **(F)** Effect of PG remodelling by plasmid-produced PBP5 on SDS sensitivity in CS703-1 and CS703-1Δ*lpp*. Ectopic PBP5 production from p*dacA* partially restores OM integrity. **(G)** Model for spatial coupling of PG biosynthesis and BAM activity in the cell. BAM-mediated OMP insertion is dampened by mature PG, resulting in OMP biogenesis being largely absent at the old poles of cells and occurring predominantly at PG growth sites.

## Discussion

Our data reveal that although BAM is uniformly distributed in *E. coli* its ability to insert OMPs into the OM is modulated by the identity of the PG immediately beneath it, an effect no doubt enhanced by the lack of mobility of OMPs in the membrane^45^. This modulation has two important physiological consequences for the bacterium. First, it synchronises insertion of OMPs with the growth and maturation of the cell wall. Such synchrony likely includes growth of the OM itself since the OMP LptD, which deposits lipopolysaccharides into the outer leaflet of the membrane, is a BamA substrate^46^. Second, biasing OMP insertion to septa/mid-cell regions enables the organism to turnover its OMPs simply by division (binary partitioning). As result, the organism can rapidly alter its OMP composition in response to a changing environment without the need for active protein degradation^45^.

## Acknowledgements

This work was funded by the European Union’s Horizon 2020 research and innovation programme under the Marie Skłodowska-Curie grant agreement No 721484 (International Training Network Train2Target) awarded to WV and European Research Council Advanced grant (742555; OMPorg), Wellcome Trust collaborative award (201505/Z/16/Z) and BBSRC UK project grant (BB/P009948/1) to CK. GM was supported by an EMBO Long-Term Fellowship (ALTF 485-2017). We thank Steven T. Rutherford, Jian Payandeh and Kelly M. Storek (Genentech Inc) for helpful discussions and comments on the manuscript, Nicholas G. Housden (Oxford) for plasmid pNGH206, Emma Elliston for providing the *fepA* expressing construct, Dr Harris Bernstein (National Institute of Diabetes and Digestive and Kidney Diseases, Bethesda, USA) for plasmid pJH114, Dr Manjula Reddy (Centre for Cellular and Molecular Biology, Hyderabad, India) for plasmid pMN86, Dr Jean-Francois Collet (de Duve Institute, Brussels, Belgium) for antibodies against Lpp and Bam proteins, Dr Christian Otten (Centre for Bacterial Cell Biology, Newcastle University) for strain CS703-1Δlpp and Sandip Kumar (Oxford) for help with ONI nanoimager experiments. We are also indebted to Nadia Halidi and the Micron Advanced Bioimaging Unit (Wellcome Strategic Awards 091911/B/10/Z and 107457/Z/15/Z) for their support & assistance in this work.

## Data availability

The data supporting the findings of the study are available in the article or available upon request from the corresponding author.

## Author contributions

G.M., F.C., C.K., and W.V. designed research; G.M. and F.C. performed research; R.C.K., D.S., T.J.K., and P.S. contributed new reagents and proteins; G.M. and F.C. analyzed data; and G.M., F.C., C.K, and W.V. wrote the paper.

## Methods

### Strains plasmids and Oligonucleotides used in this study

The bacterial strains, plasmids and oligonucleotides used in this study are provided in Table S1, Table S2 and Table S3, respectively.

### Construction of pNGH206

For the construction of the pNGH206 plasmid, site-directed mutagenesis was used to introduce a solvent accessible cysteine (K469C) in the cytotoxic domain of a construct in which the N-terminal 62 amino acids of the colicin had been deleted (Δ^2-61^ ColE9).

### Construction of pBAD33-*dacA*

The *dacA* gene was amplified by polymerase chain reaction (PCR) with Q5 Polymerase (NEB), using chromosomal DNA from *E. coli* BW25113 as template and the oligonucleotides dacA_F and dacA_R (Table S3). Plasmid pBAD33 was amplified by PCR with Q5 Polymerase using oligonucleotides pBAD33_FR and pBAD33_RF (Table S3). Insert and vector were joined by ligation-independent cloning^48^. Positive clones were selected on LB agar + 25 μg/ml chloramphenicol and identified by colony PCR using GoTaq G2 Polymerase (Promega). The correct sequence of the cloned *dacA* gene was confirmed by double-strand sequencing using specific oligonucleotides pBAD33_seq_F1, dacA_seq_R1, dacA_seq_F2 and pBAD33_seq_R2 (Table S3). All oligonucleotides were obtained from Eurogentec (Seraing, Belgium).

### Expression and purification of antibodies and colicins

Antibodies and engineered colicins used are listed in table S4. **αBamA MAB2 Fabs:** A construct suitable for periplasmic expression of Fab in *E. coli* and containing a sequence coding for Fab fragments of MAB2 was cloned; It was transformed into 34B8 *E. coli* cells and expressed at 30°C under control of the *phoA* promoter in CRAP phosphate-limiting autoinduction medium (PMID: 12009210) supplemented with carbenicillin (50 μg/mL). After 24 hr, cells were harvested and resuspended in PBS supplemented with one complete EDTA-free Protease Inhibitor Cocktail tablet (Roche) per 50 mL of lysis buffer, lysozyme (0.125 mg/mL), and benzonase (0.01 mg/mL). The prepared suspension was microfluidized at 15,000 psi and clarified at 50,000 x g for 30 min at 4°C. The supernatant was then resolved on protein G Sepharose beads equilibrated with PBS, using 2 mL packed resin volume per original gram of cell paste. The column was washed extensively with PBS and Fabs were eluted under mildly acidic conditions (0.56 % glacial acetic acid pH 3.6). Eluted Fabs were immediately dialyzed overnight at 4°C against buffer containing 500 mM NaCl, 10% glycerol, and 100 mM Tris (pH 8.0). Fabs were further purified on an S75 16/60 gel filtration column (GE Healthcare) using PBS (pH 7.2) as the running buffer.MAB2 Fab fragments of were labelled by using Alexa Fluor 488 Protein Labeling Kit (Thermo Fisher Scientific) following the manufacturer’s instructions. The fluorescently labelled Fabs were passed over HiPrep desalting column (GE Healthcare) to remove the excess dye. Peak fractions were collected and concentrated, and the degree of labelling (=1.42) determined using liquid chromatography mass spectrometry (LC–MS). **ColicinE9-AF488 expression and purification:** The expression and purification of this protein has been previously described^8^. Here, we used a modified construct with a single cysteine (Δ^2-61^ColE9 K469C-Im9_His6_). Cys469 in the C-terminal DNase domain of these ColE9 constructs was labelled with a 3-fold Alexa Fluor 488-maleimide (Invitrogen) as previously described^8^. The labelling efficiency (typically 0.8 fluorophores/protein) was estimated spectrophotometrically (V550 spectrophotometer, Jasco). **Colicin B-GFP/mCherry expression and purification:** The expression and purification of these proteins has been previously described at Cohen-Khait et *al*., 2021^26^.

### Expression and purification of soluble constructs of BamA P1,2 (residues 21-174), P3,4 (residues 175-345), P4,5 (266-422), BamB, BamC and BamE

Constructs for BamB and BamE were synthesised without their periplasmic export sequences with the cysteine at the beginning of the mature protein mutated to serine to remove their N-terminal acylation sites and cloned into pET22b(+) (Genscript, Novagen). A construct for BamC was synthesised lacking its periplasmic export sequence and N-terminal acylation site (residues 26–344) and incorporated into the pET16b expression vector with an N-terminal 6xHis-tag (Genscript, Novagen)^49^. A construct for BamA P1,2 (residues 21-174) was cloned into pQE70, with a C-terminal 4xHis-tag^50^. Constructs for P3,4 (residues 175-345) and P4,5 (residues 266-422) were synthesised and cloned into pET26b(+), with a C-terminal 6xHis-tag (Genscript, Novagen).

Cells containing the appropriate plasmid were grown in LB media supplemented with 100 µg/ml ampicillin for BamB, BamC, BamE and P1,2, and 30 µg/ml kanamycin for P3,4 and P4,5, to an OD_600_ of 0.4 and protein expression induced by the addition of 1 mM IPTG, at 18°C overnight. Cultures were harvested by centrifugation (6000×*g*, 15 min), resuspended in 50 mM sodium phosphate pH 7.5, 300 mM NaCl, 10 mM imidazole with EDTA-Free protease inhibitor tablets (Roche) and lysed using an Emulsiflex C3 cell disruptor (Avestin). The lysate was centrifuged at 75000×*g* for 45 minutes at 4°C to pellet insoluble material. The supernatant was filtered through a 0.45 µM filter (Millipore), then purified via immobilised metal affinity chromatography using a 5 mL His-trap HP column (GE Healthcare) in sodium phosphate buffer pH 7.5 followed by size exclusion chromatography in 50 mM Sodium phosphate pH 7.5, 300 mM NaCl, using a Superdex 75 26/60 column (GE Healthcare). Fractions were assessed by SDS-PAGE, combined, concentrated using an Amicon Ultra 10 kDa MWCO centrifugal concentrator (Millipore) and stored at 4°C for immediate use or frozen in liquid nitrogen and stored at -80°C.

### Expression and purification of BamCD and BamCDE

Plasmids expressing BamCD (pSK46 – Hagan *et al*., 2010)^39^ and BamE with a C-terminal 6xHis-tag (pBamE-His) ^39^ were transformed separately into BL21(DE3) cells (New England Biolabs). The cells were grown in LB broth (supplemented with 50 µg/ml streptomycin for BamCD and 100 µg/ml ampicillin for BamE), at 37°C, to an OD_600_ of 0.4 and protein expression induced by the addition of 0.5 mM IPTG, overnight, at 18°C. Cells were harvested separately by centrifugation (6000×*g*, 15 min), resuspended in 20 mM Tris pH 8.0, 150 mM NaCl with EDTA-Free protease inhibitor tablets (Roche) and lysed separately, using an Emulsiflex C3 cell disruptor (Avestin). The lysates were spun separately at 10000×*g* for 30 min at 4°C and the supernatant centrifuged at 100000×*g* for 45 minutes to harvest membranes. The membranes were solubilised (1 ml of buffer for every 40 mg of membrane) with 50 mM Tris pH 8.0, 150 mM NaCl, 1% DDM (Anatrace), combined and then rotated at 4°C for two hours. The solubilised membranes were centrifuged at 50000×*g* for 30 min; the supernatant filtered through a 0.45 µm filter (Millipore), and then bound to equilibrated Ni-NTA agarose beads (Qiagen) overnight at 4°C. The beads were washed with three column volumes of 50 mM Tris pH 8.0, 150 mM NaCl, 50 mM Imidazole, 0.03% DDM and the protein eluted with two column volumes of 50 mM Tris pH 8.0, 150 mM NaCl, 500 mM imidazole, 0.03% DDM. Fractions were assessed by SDS-PAGE and those containing BamCDE were pooled and further purified through size exclusion chromatography using a Superdex 200 16/600 column (GE Healthcare) in 50 mM Tris pH 8.0, 150 mM NaCl, 0.03% DDM. Fractions were further assessed by SDS-PAGE, combined and stored at 4°C for immediate use or frozen in liquid nitrogen and stored at -80°C.

BamCD were co-purified from a modified version of plasmid pSK46 carrying a 6xHis-tag at the C-terminus of BamC following the same protocol, omitting the steps with plasmid pBamE-His.

### Purification of BamABCDE

The protocol was adapted from previous reports^40, 41^. Plasmid pJH114^40^ was transformed into *E. coli* BL21(DE3). Cells were grown in LB (10 g/L NaCl) + 100 µg/ml ampicillin up to OD_600_ ∼0.5-0.6, then protein overproduction was induced with 0.4 mM IPTG by incubating at 37°C for 90 min with orbital shaking (175 rpm). Cells were harvested (6200×*g*, 4°C, 15 min), resuspended in Buffer A (20 mM Tris/HCl, pH 8.0) and disrupted by sonication. Membranes were harvested by ultracentrifugation (130000×*g*, 4°C, 1 h) and solubilised in Buffer B containing 50 mM Tris/HCl, 150 mM NaCl, 1% N-dodecyl β-D-maltoside (DDM, Avanti) at pH 8.0 by incubating for 1 h on ice. The sample was incubated with 2 ml per liter of culture volume of Ni-NTA agarose beads (Qiagen) and rotated overnight at 4°C on a tube roller. Beads were washed in Buffer C (50 mM Tris/HCl, 150 mM NaCl, 50 mM imidazole, 0.05% DDM, pH 8.0) and proteins eluted in Buffer D (50 mM Tris/HCl, 150 mM NaCl, 500 mM imidazole, 0.05% DDM, pH 8.0). Eluted fractions were concentrated to ∼500 µl in ultrafiltration units and applied to a Superdex 200 (10/300) column (GE Healthcare), in filtered and degassed Buffer E (50 mM Tris/HCl, 150 mM NaCl, 0.05% DDM, pH 8.0) at 0.5 ml/min, collecting 500 µl fractions. Protein purity and yield was analysed by SDS-PAGE. Fractions containing BamABCDE were combined and immediately reconstituted into proteoliposomes or snap-frozen in liquid nitrogen and stored in small aliquots at -80°C.

### Purification of SurA

The protocol was adapted from previous reports^39, 40^. SurA was overproduced in *E. coli* BL21(DE3) by growing cells in LB (10 g/L NaCl) + 50 µg/ml kanamycin up to an OD_600_ of ∼1.0. The temperature was shifted to 16°C and 0.1 mM (final conc.) IPTG was added and the cells were incubated at 16°C for ∼16-18 h. Cells were harvested (6200×*g*, 4°C, 15 min), resuspended in Buffer A (20 mM Tris/HCl, pH 8.0) and disrupted by sonication. The soluble fraction was incubated with 2 ml per liter of culture volume of Ni-NTA agarose beads (Qiagen) and rotated overnight at 4°C on a tube roller. Beads were washed in Buffer B (20 mM Tris/HCl, 50 mM imidazole, pH 8.0) and the protein eluted in Buffer C (20 mM Tris/HCl, 500 mM imidazole, pH 8.0). Eluted fractions were dialysed against Buffer D (20 mM Tris/HCl, 10% glycerol) overnight at 4°C, then concentrated to ∼5 ml and applied to a Superdex 75 (16/600) column (GE Healthcare), in filtered and degassed Buffer D at 1 ml/min. Eluted fractions were analysed by SDS-PAGE to assess protein purity and yield. Fractions containing SurA were combined and concentrated to 250-300 µM in a Vivaspin Turbo 10 kDa centrifugal concentrator (Sartorius), and stored in aliquots at -80°C.

### Purification of OmpT

An adapted protocol was used ^41^. OmpT was overproduced as cytoplasmic inclusion bodies in *E. coli* BL21(DE3) by growing cells in LB (10 g/L NaCl) + 50 µg/ml kanamycin up to OD_600_ ∼0.5- 0.6, adding 1 mM IPTG and incubating for 4 h at 37°C. Cells were harvested (6200×*g*, 4°C, 15 min), resuspended in Buffer A (50 mM Tris/HCl, 5 mM EDTA, pH 8.0) and disrupted by sonication. The insoluble fraction was collected by centrifugation (4500×*g*, 4°C, 15 min) and resuspended in Buffer B (50 mM Tris/HCl, 2% Triton X-100, pH 8.0), then incubated for 1 h at room temperature with gentle shaking. Inclusion bodies were pelleted (4500×*g*, 4°C, 15 min) and washed twice in Buffer C (50 mM Tris/HCl, pH 8.0) by incubating for 1 h at room temperature, then solubilised in Buffer D (25 mM Tris/HCl, 6 M guanidine-HCl, pH 8.0). The supernatant was filtered, concentrated to ∼5 ml in a Vivaspin Turbo 10 kDa centrifugal concentrator (Sartorius), and applied to a Superdex 75 (26/600) column (GE Healthcare) with filtered and degassed Buffer D at 1 ml/min. Eluted fractions were analysed by SDS-PAGE to assess protein purity and yield. Fractions containing OmpT were combined and stored in aliquots at -80°C.

### Purification of MepM

An adapted protocol was used^32^. Soluble MepM carrying a C-terminal 6×His-Tag was overproduced in the cytoplasm of *E. coli* BL21 (DE3) from plasmid pMN86. Cells were grown in LB (10 g/L NaCl) + 100 µg/ml ampicillin at 37°C with shaking up to OD_600_ ∼0.6. The culture was shifted to 25°C and supplemented with 50 μM IPTG after 30 min to induce protein overproduction. Induction was followed for 2 h. Cells were harvested by centrifugation (6200×*g*, 4°C, 15 min) and resuspended in Buffer A (25 mM Tris/HCl, 300 mM NaCl, 10 mM MgCl_2_, 20 mM imidazole, 10% glycerol, pH 7.0). Cells were disrupted by sonication and the soluble fraction applied to a 5 ml-HisTrap HP column at 1 ml/min. The column was washed with 5 column volumes of Buffer A at 1 ml/min. MepM was eluted at 1 ml/min in Buffer B (25 mM Tris/HCl, 300 mM NaCl, 10 mM MgCl_2_, 400 mM imidazole, 10% glycerol, pH 7.0). Fractions containing MepM were pooled and dialysed overnight at 4°C against Buffer C (25 mM Tris/HCl, 300 mM NaCl, 10 mM MgCl_2_, 10% glycerol, pH 7.0), then concentrated to a volume of ∼5 ml and applied to a Superdex 75 (16/600) column (GE Healthcare), in filtered and degassed Buffer C at 1 ml/min. Eluted fractions were analysed by SDS-PAGE and fractions containing MepM were combined and stored in aliquots at -80°C.

### Isolation of PG sacculi

An adapted protocol was used^51^. Cells were grown in 4 L of LB (10 g/L NaCl) at 37°C with orbital shaking (175 rpm), up to OD_600_ ∼0.5-0.6. Cultures were incubated on ice for 10 min to stop cell growth. Cells were harvested (6200×*g*, 4°C, 15 min) and resuspended in 40 ml of ice-cold Milli-Q water. The cell suspension was added drop-wise to 40 ml of boiling 8% SDS and boiled with vigorous stirring for 30 min. After cooling down to room temperature, sacculi were collected by ultracentrifugation (130000×*g*, 25°C, 1 h) and washed in Milli-Q water. Ultracentrifugation and washing steps were repeated until samples were SDS-free^51^. Sacculi were resuspended in 9 ml of 10 mM Tris/HCl, 10 mM NaCl, pH 7.0, supplemented with 1.5 mg of α-amylase (Sigma-Aldrich) and incubated at 37°C for 2 h. Samples were supplemented with 2 mg of Pronase E (Sigma-Aldrich) and incubated at 60°C for 1 h. Reactions were stopped by adding 4% SDS (1:1 v/v) and boiling at 100°C for 15 min, then samples washed until SDS-free as before. Purified sacculi were resuspended at ∼10 mg/ml in 0.02% NaN_3_ and stored at 4°C.

### Preparation of disaccharide tetrapeptide chains (Tetra^n^) for MST experiments

Isolated sacculi from *E. coli* BW25113Δ6LDT^29^ were incubated at 5.6 mg/ml with MepM (3 µM) in 25 mM Tris/HCl, 150 mM NaCl, 0.05% Triton X-100, pH 7.5. Negative control samples containing no sacculi (mock digests) for MST experiments were prepared in parallel. Samples were incubated for ∼18 h at 37°C with shaking and boiled at 100°C for 10 min. Soluble PG fragments were collected by centrifugation (17000×*g*, room temperature, 15 min) and dialysed against 50 mM sodium phosphate, 150 mM NaCl, pH 7.0 for ∼24 h at room temperature in a 3-5 kDa dialysis membrane. Dialysed DS-tetra chains were stored at -20°C.

### HPLC analysis of PG and Tetra^n^

Purified PG sacculi or Tetra^n^ (∼100 µg) were digested with cellosyl (0.5 μg/ml) for 16-18 h at 37°C in 20 mM sodium phosphate, pH 4.8 with shaking (1000 rpm). Digestions were stopped by boiling at 100°C for 10 min. Muropeptides were collected after centrifugation (15000×*g*, 15 min), reduced with sodium borohydride as described^51^ and analysed by reversed-phase HPLC at 55°C in a 90- or 180-min linear gradient from 50 mM sodium phosphate, pH 4.31 to 75 mM sodium phosphate, pH 4.95, 15% methanol, on an Agilent 1220 Infinity HPLC system (Agilent). Relative concentrations of muropeptides from different sacculi preparations were estimated by comparing the total peak area from the integration of the UV signal from HPLC chromatograms^51^.

### PG pulldown assay

The protocol was adapted from previous reports^30, 31^. PG sacculi (∼1 mg) were incubated with purified protein (5 µM) in PG binding buffer (50 mM Tris/maleate, 50 mM NaCl, 10 mM MgCl_2_, pH 7.5) in a total volume of 100 µl. Samples were incubated on ice for 30 min, then pelleted by centrifugation (17000×*g*, room temperature, 10 min) and the supernatant was collected (supernatant fraction, S). PG pellets were washed by resuspending in 200 µl of PG binding buffer and pelleting again (17000×*g*, room temperature, 10 min) and the supernatant was recovered (wash fraction, W). PG-bound protein was released from sacculi by resuspending the PG pellet in 100 µl of 2% SDS and stirring for 1 h at room temperature. Samples were centrifuged (17000×*g*, room temperature, 10 min) and the supernatant was collected (pellet fraction, P). Fractions S, W and P were analysed by SDS-PAGE on 15% polyacrylamide gels and proteins visualised by Coomassie Blue staining. Protein retention in fraction P indicated binding of the protein to PG sacculi. For experiments performed with the full-length BamABCDE complex, PG binding buffer was supplemented with 0.05% Triton X-100. PG pull-down experiments with SurA were performed in 20 mM Tris/HCl, pH 6.5.

### Microscale thermophoresis (MST)

Purified proteins were labelled with Red-NHS (NanoTemper Technologies) in MST labelling buffer (50 mM sodium phosphate, 150 mM NaCl, 10% glycerol, pH 7.0) according to the manufacturer’s protocol. The concentration of fluorescently-labelled proteins and efficiency of labelling was determined spectrophotometrically. MST experiments were performed as follows: serial dilutions of Tetra^n^ (from ∼5.6 mg/ml to ∼0.2 μg/ml, 16 samples in total) or mock digestions were prepared in MST buffer in a total volume of 10 μl, mixed to an equal volume of labelled protein at 100 nM in MST buffer supplemented with 0.1% reduced Triton X-100 (Sigma-Aldrich) in order to obtain a serial dilution of ligand from ∼2.8 mg/ml to ∼0.1 μg/ml, a final protein concentration of 50 nM, detergent concentration of 0.05% and reaction volume of 20 μl. Samples were incubated for 5 min on ice and 5 min at room temperature and loaded into standard-coated MST capillaries. Measurements were performed in a Monolith NT.115 (NanoTemper Technologies). LED power for each set of experiment was chosen in order to obtain initial fluorescence values between 200 and 2500 counts for each individual protein during the capillary scan. Thermophoresis was analysed at the steady-state region of each thermogram. Curve fitting for *K*_D_ measurements was performed by plotting the normalised fluorescence intensity (F_norm_) at the steady state for each sample against ligand concentration according to a 1:1 binding model, as an average of three independent experiments. Results were analysed using the MO-Affinity Analysis software (NanoTemper Technologies).

Alternatively, for proteins which exhibited variations in initial fluorescence greater than ± 10% of the average fluorescence along the serial dilution prior to the application of the temperature gradient, curve fitting was performed by directly plotting the initial fluorescence of samples against ligand concentration. *K*_D_ was calculated assuming a 1:1 binding model from an average of three independent experiments. To confirm that initial fluorescence changes were ligand-dependent, SDS-denaturation (SD) tests were performed as follows: after preparing the serial dilution as for the main interaction experiments, the initial fluorescence of three capillary samples representative of the bound fraction (capillary 1, 2 and 3) and three capillary samples representative of the unbound fraction (capillary 14, 15 and 16) was first measured to determine the differences between bound and unbound states. Samples were then centrifuges at 15000×*g* for 10 minutes, and the supernatant mixed 1:1 with 2× SD-mix (40 mM DTT, 4% SDS) and boiled at 95°C for 10 minutes. The initial fluorescence for the chosen samples was then measured again and compared to the initial fluorescence observed before the SD-tests. The initial difference was confirmed to be ligand-dependent if initial differences in fluorescence between samples from the bound and unbound fractions disappeared by the SDS-treatment. SD-tests were performed in triplicate and analysed using the MO-Affinity Analysis software (NanoTemper Technologies).

### Reconstitution of BamABCDE into liposomes

The protocol was adapted from previous reports^39, 40^. *E. coli* polar lipids (Avanti) were resuspended at 20 mg/ml in water, well dispersed by sonication and 200 µl were mixed with 1 ml of freshly purified BAM complex, and incubated on ice for 5 min. The mixture was diluted with 20 ml of 20 mM Tris/HCl, pH 8.0 and incubated on ice for 30 min. Proteoliposomes were pelleted by ultracentrifugation (135000×*g*, 4°C, 30 min) and washed in 20 ml of 20 mM Tris/HCl pH 8.0, pelleted again and resuspended in 800 μl of 20 mM Tris/HCl pH 8.0. Efficiency of reconstitution was assessed by SDS-PAGE, analysing the supernatant, wash and pellet fractions. Aliquots (20 μl) of proteoliposomes were snap-frozen in liquid nitrogen and stored at -80°C.

### *In vitro* BAM activity assay

Two 25 µl sub-reactions (A and B) were assembled in 20 mM Tris-HCl, pH 6.5 as follows: sub-reaction A contained SurA (140 µM) and OmpT (20 µM); sub-reaction B contained BAM proteoliposomes (2 µM), fluorogenic peptide (Peptide Synthetics) (2 mM). The two sub-reactions were assembled in half-area, black microplates (Corning) and incubated at 30°C for 5 min, and mixed to start OmpT folding (final concentrations: 1 µM BAM proteoliposomes, 1 mM fluorogenic peptide, 70 µM SurA and 10 µM OmpT in a total volume of 50 µl). When required, PG sacculi or Tetra^n^ prepared in 20 mM Tris-HCl, pH 6.5 were supplemented to sub-reaction B. Fluorescence emission (excitation at 330 nm, emission at 430 nm) upon cleavage of the fluorogenic peptide by folded OmpT was monitored at 30°C for 1 h 20 min after in a FLUOStar Microplate Reader (BMG Labtech), with readings every 20 s and 5 s orbital shaking prior to each reading. Three independent replicates were analysed for each experiment. Activity rates for each replicate were analysed over the linear range in the fluorescence release curve, averaged and converted into percentage relative to control reactions containing no PG.

### *In vitro* BAM activity assay with pre-folded OmpT

For experiments in which OmpT was folded prior to fluorescence measurements, reactions were assembled as described and incubated for 2 h 30 min at 30°C with orbital shaking to allow BAM-mediated OmpT assembly. Samples were then mixed with 50 µl reactions containing either 2 mM fluorogenic peptide or 2 mM fluorogenic peptide and 5 mg/ml sacculi from *E. coli* MC1061 in 20 mM Tris-HCl, pH 6.5, and OmpT activity was monitored as described.

### *In vitro* BAM activity assay with an excess of SurA

Experiments containing an excess of SurA were performed as follows. SurA (15 µM) was mixed with 1 mg of sacculi from *E. coli* MC1061 in 20 mM Tris-HCl, pH 6.5, in a total volume of 200 µl. Control samples contained no PG. Samples were incubated on ice for 30 min, then split in half. One half was added to OmpT folding reactions, assembled as described (BAM activity control reactions containing no PG and no extra SurA, or PG only, were included). The other half was used to monitor PG binding of SurA as described. OmpT activity was measured as above.

### Western Blot analysis

*E. coli* cell suspensions were mixed 1:1 with 2× SDS-PAGE loading buffer (100 mM Tris-HCl, pH 6.8, 4% SDS, 0.2% bromophenol blue, 20% glycerol, 10% β-mercaptoethanol) and boiled at 100°C for 10. Samples were loaded on 15% polyacrylamide gels and proteins resolved by SDS-PAGE, then transferred to nitrocellulose membranes and probed with specific primary antibodies (α-BamA 1:40000; α-BamB 1:3000; α-BamC 1:20000; α-BamE 1:1500; α-Pal 1:2500; α-CpoB 1:2500; α-PBP5 1:1000; α-Lpp 1:3000). Goat anti-rabbit HRP-IgG (Sigma Aldrich, 1:5000) were used as secondary antibodies. Western Blots were developed using ECL™ Prime Western Blotting System (GE Healthcare).

### *In vivo* cross-linking of Bam proteins to PG

The protocol was adapted from a previous report^33^. Briefly, *E. coli* MC1061 was grown in 50 ml of LB (5 g/L NaCl) at 37°C by orbital shaking up to OD_600_ ∼0.5. Cells were pelleted (4500×*g*, room temperature, 10 min), the cell pellet washed with 50 ml of phosphate-buffered saline (PBS) three times and the OD_600_ adjusted to 2.0 with PBS. 3,3’-dithiobis (sulfosuccinimidyl propionate) (DTSSP, ThermoFisher) was freshly dissolved in 5 mM sodium citrate, pH 5.0 and added to cells to a final concentration of 0.5 mM. Cells were incubated at room temperature for 10 min. The cross-linking reaction was quenched by adding Tris/HCl, pH 8.0 to a final concentration of 50 mM, incubating at room temperature for 15 min. Whole cell samples were taken for Western Blot analysis by concentrating 300 µl of cells 3-fold and mixing 1:1 v/v with SDS-PAGE buffer (200 mM Tris/HCl pH 6.8, 4% SDS, 20% glycerol, 0.2% bromophenol blue) with or without 10% β-mercaptoethanol. The rest of the bacterial suspension was added drop-wise to an equal volume of boiling 8% SDS and boiled with vigorous stirring for 30 min to isolate PG sacculi. After cooling down to room temperature, sacculi were pelleted by ultracentrifugation (130000×*g*, room temperature, 1 h) and washed twice in 2% SDS, then resuspended in ∼100 μl of 2% SDS. To analyse PG-bound proteins, sacculi suspensions were boiled in SDS-PAGE buffer with or without 10% β-mercaptoethanol at 100°C for 10 min, briefly centrifuged, and supernatants loaded on 15% polyacrylamide gels, together with whole cell samples taken after cross-linking. Proteins were transferred to nitrocellulose membranes for Western Blot analysis.

### SDS sensitivity assay (Spot-plate)

*E. coli* strains were grown from a single colony in LB + 25 µg/ml chloramphenicol at 37°C by orbital shaking for ∼16-18 h. The OD_600_ was adjusted to 2.0 and cells were serially 10× diluted to 10^-7^ in growth medium, then plated with a pin replicator on LB agar + 25 µg/ml chloramphenicol + 0.2% arabinose, with or without 2% SDS. Plates were incubated at 37°C and photographed after 24 h of incubation.

### Cell preparation for live microscopy

Overnight LB (10 g/litre tryptone, 10 g/litre NaCl, 5g/later yeast extract [pH 7.2]), supplemented M9-glucose media (0.4% (w/v) D-glucose, 2 mM MgSO_4_, 0.1 mM CaCl_2_, 1mg/ml NH_4_Cl, 0.05% (w/v) casamino acids) cultures were grown at 37 °C and diluted 1:100 into fresh medium with appropriate antibiotics. Cultures were grown at 37 °C, unless stated otherwise, to mid-log phase (OD_600_ = 0.2-0.7) and cells were centrifuged at 7000 × g for 1 min. For translation inhibition experiments, cells were treated with chloramphenicol (30 µg ml^−1^) 30 minutes before samples were taken. Agar pads were prepared by mixing supplemented M9- glucose medium or PBS with 1% agarose and pouring 150 µl into 1.5 × 1.6 gene frame (Thermo Scientific #AB0577) attached to the slide. For pad formation, the gene frame was sealed by a coverslip until agarose solidified. Six microliters of cells were pipetted onto the agar pad, allowed to dry and sealed with a clean coverslip. For the induction of OMPs from a plasmid, 0.4% (w/v) arabinose was added directly into the growing culture 7 minutes before samples were taken, unless stated otherwise.

For live cells labelling an equivalent of 1ml of cells at OD_600_ = 0.25 were pelleted by centrifugation (7000 x g, 1 min) and the samples were resuspended in supplemented M9- glucose media containing 200nM fluorescently labelled MAB2. The labelling was carried out for 20 min at room temperature with mixing by rotary inversion in an opaque tube. Subsequently the cells were washed twice (M9-glucose) by pelleting (7000 x g, 1 min) and finally resuspended in ∼50 μl M9-glucose. For fixed cells labelling, a similar amount of cells were pelleted by centrifugation (7000 x g, 1 min) and the samples were resuspended in 4% formaldehyde (in PBS) at 4°C immediately after sampling. After 20 minutes, cells were pelleted by centrifugation (7000*g*, 1 min) and resuspended in 100 μl of fresh supplemented PBS containing 200nM fluorescently labelled Fabs or colicins. Labelling was carried out as with live cells. After labelling, the cells were washed thrice (in PBS) by pelleting (7000 x g, 1 min) and finally resuspended in ∼50 μl PBS. To improve binding of the MAB2 Fabs in co-labelling experiments, after the labelling step cells were washed once (in PBS), pelleted (7000g, 1 min) and resuspended in 4% formaldehyde (in PBS) at 4°C for further 20 minutes. Cells were then washed twice (PBS) and resuspended in ∼50 μl PBS.

The Fluorescent D-Amino Acid (FDAA) 7-hydroxycoumarincarbonylamino-D-alanine (HADA) (Tocris Bioscience) was used for cell wall labelling. The labelling was carried as described previously^53^ with minor adjustments. The final concentration of HADA in the growing culture was 500µM and the incubation time varied according to the aims of the experiment. When both PG and OMP labelling were included, the HADA labelling protocol was used first. After completing the last step of HADA labelling (fixation), samples were labelled using the relevant colicins as described above.

### TIRFM acquisition

Live cells were imaged using an Oxford NanoImager (ONI) superresolution microscope equipped with four laser lines (405/473/561/640 nm) and ×100 oil immersion objective (Olympus 1.49 NA). Fluorescence images were acquired by scanning a 50 µm × 80 µm area with a 473 nm laser for AF488 & GFP labelled proteins (laser power 1.4-2.3 mW) or 561 nm for mCherry labelled proteins (laser power 2.1-3.4 mW). The laser was set at 50° incidence angle (200 ms exposition), resulting in a 512 × 1024 pixel image. Images were recorded by NimOS software associated with the ONI instrument. Each image was acquired as a 20-frame stack for brightfield and fluorescence channels, respectively. For analysis, images were stacked into composite images using average intensity as a projection type in ImageJ (version 1.52p). To ensure non-uniform fluorescence of labelled OMPs was not the result of proximity to the coverslip, equivalent images were taken in epifluorescence and by 3D-SIM microscopy where such potential bias was absent (see below).

### Epifluorescence and SIM Imaging

Cells were imaged using Deltavision OMX V3 Blaze microscopy system (GE) equipped with four laser lines (405/488/594/633 nm), a standard or a Green/Red drawer filter set and a ×60 oil immersion objective (Olympus 1.42 NA). For both conventional and SIM imaging 1.512 index refraction immersion oil was used for AF488 & GFP labelled proteins and 1.514 index refraction immersion oil was used for AF488/mCherry dual-color imaging. Conventional fluorescence images were acquired by imaging a 42 μm x 42 μm area with the 488 nm laser (5.7 mW, 500ms exposure) resulting in a 512 x 512 pixel image. For SIM acquisition, a similar area was imaged using the 488 nm laser (2.7 mW, 200 ms exposure). Image stacks of 1-1.5μm thickness were taken with 0.125 μm z-steps and 15 images (three angles and five phases per angle) per z-section and a 3D-structured illumination with stripe separation of 213 nm and 238 nm at 488 nm and 594 nm, respectively. The SIMcheck plugin (ImageJ) was used to assess the data quality of SIM images. Image stacks were reconstructed using Deltavision softWoRx 7.2.0 software with a Wiener filter of 0.003 using wavelength specific experimentally determined OTF functions. Average intensity and 3D projections of 3D-SIM images were generated using ImageJ (V1.52p)

For the acquisition of multi-channel images, a DIC image was taken first followed by an imaging sequence which minimized any possible overlap between channels. Fluorophores with higher excitation/emission spectra were imaged first and the fluorescent signal was bleached prior to the acquisition of the following channel. Alignment of dual-colour images was carried out using TetraSpeck™ Microspheres, 0.1 µm (ThermoFisher scientific) and the channel aligner tool (ImageJ V1.52p).

### Image and data analysis

2D-SIM images of BamA labelled cells were binarized and Regions Of Interest (ROI’s) were generated. Non-distinct islands were manually excluded. The size of each island was calculated based on its Ferret’s diameter (ImageJ V1.52p). For measurement of septal cell widths, the DIC and epifluorescence images were overlaid and the HADA channel used to determine the location of the developing septum along the long axis of the cell. Measuring the width of the cell at the selected region was done using the DIC channel (ImageJ V1.52p).

For the detection of islands and creating integrated localisation maps an Unsharp mask filter (Radius 2px, Mask weight 0.5) was applied to the raw images and BamA islands were identified using the Find Maxima process (Prominence > 600) (imageJ) and ROI’s were created. The fluorescence intensity of each ROI was measured using the raw images and background fluorescence was subtracted. Integrated localization maps of BamA islands were created using the ImageJ plugin MicrobeJ (v5.13m)^54^. For the detection of BamA islands with microbeJ, Unsharp Mask filter was applied as described above. Cells and maxima points detection was carried out using plugin and the auto segmentation was manually corrected to exclude from the analysis improperly detected or clustered cells.

Measurement of individual fluorescent intensity profiles and calculation of the normalized fluorescence distribution of OMPs was carried out as follows: For the early experiments (Figure 1, and the related supplementary figures) fluorescence distribution across the long axis of the OM was determined by the plot profile function (ImageJ V1.52). After measuring the raw values, they were normalized to 0−1 scale for comparison between cells. To enable the integration of fluorescence intensity distribution from cells of different lengths, the long axis of each cell was normalized to 0−100. In later experiments (Figures 2-4, and the related extended data figures) the measurement of the fluorescence intensity profile and normalization of the long axis between 0-100 was automated using the MicrobeJ plugin (v5.13m). After the integration of profiles from all cells, the value at the poles was set to 1 and the rest of the values were normalized accordingly. Normalization was done using Excel and the data was plotted in GraphPad Prism 8 software.

Demographs presenting the fluorescence distribution of OMPs in a large population of cells were created using the MicrobeJ plugin (v5.13m). Each demograph included cells from at least 3 different fields of view. For all experiments, cells were sorted by length in order to highlight the different cell cycle stages. For induced OMP biogenesis experiments the pole with the higher fluorescence intensity was aligned to the top in order to highlight the unipolar distribution pattern.

For Co-localisation measurements the two compared channels were overlaid (following channel alignment) using the Merge Channels tool (ImageJ V1.52) to confirm that the cells are properly aligned. Next, the fluorescence image from the 405nm channel was thresholded and a region of interest (ROI) including only cell containing regions was created. The Coloc 2 tool was used to calculate the Pearson correlation coefficient of the ROI between the different fluorescence channels. Cytofluorogram plotting was done using the imageJ plugin JACoP^55^.

## Extended Data Figures Legends

**Table S1.** *E. coli* strains used in this study.

| Strain | Features | Reference |
| --- | --- | --- |
| BW25113 | *lacI<sup>+</sup>rrnB<sub>T14</sub> ΔlacZ<sub>WJ16</sub> hsdR514*<br>*ΔaraBAD<sub>AH33</sub> ΔrhaBAD<sub>LD78</sub> rph-1*<br>*Δ(araB-D)567 Δ(rhaD-B)568*<br>*ΔlacZ4787(::rrnB-3) hsdR514 rph-1* | Datsenko & Wanner,<br>2000 |
| BW25113Δ6LDT | BW25113 *ΔldtA ΔldtB ΔldtC*<br>*ΔldtD ΔldtE ΔldtF* | Kuru *et al.*, 2017 |
| CS109 | W1485 *glnV rpoS rph* | Denome *et al.*, 1999 |
| CS12-7 | CS109 *ΔdacA* | Denome *et al.*, 1999 |
| CS703-1 | CS109 *ΔmrcA ΔdacB ΔdacA*<br>*ΔdacC ΔpbpG ΔampC ΔampH* | Meberg *et al.*, 2001 |
| CS703-1Δlpp | CS109 *ΔmrcA ΔdacB ΔdacA*<br>*ΔdacC ΔpbpG ΔampC ΔampH Δlpp* | Dr Christian Otten<br>(Newcastle<br>University) |
| GNE 4077 | BW25113 *ompT, waaD::Cm BamA_barr*<br>(*K. pneumoniae*)::Gm | Storek *et al* |
| MC1061 | K-12 F<sup>-</sup> λ<sup>-</sup> *Δ(ara-leu)7697 [araD139]B/r*<br>*Δ(codB-lacI)3 galK16 galE15 e14-*<br>*mcrA0 relA1 rpsL150(Str<sup>R</sup>) spoT1 mcrB1*<br>*hsdR2(r<sup>-</sup>m<sup>+</sup>)* | Casadaban & Cohen,<br>1980 |
| RK5016 | MC4100, *metE70, argH, btuB, recA* | Heller *et al.*, 1985 |

**Table S2.**
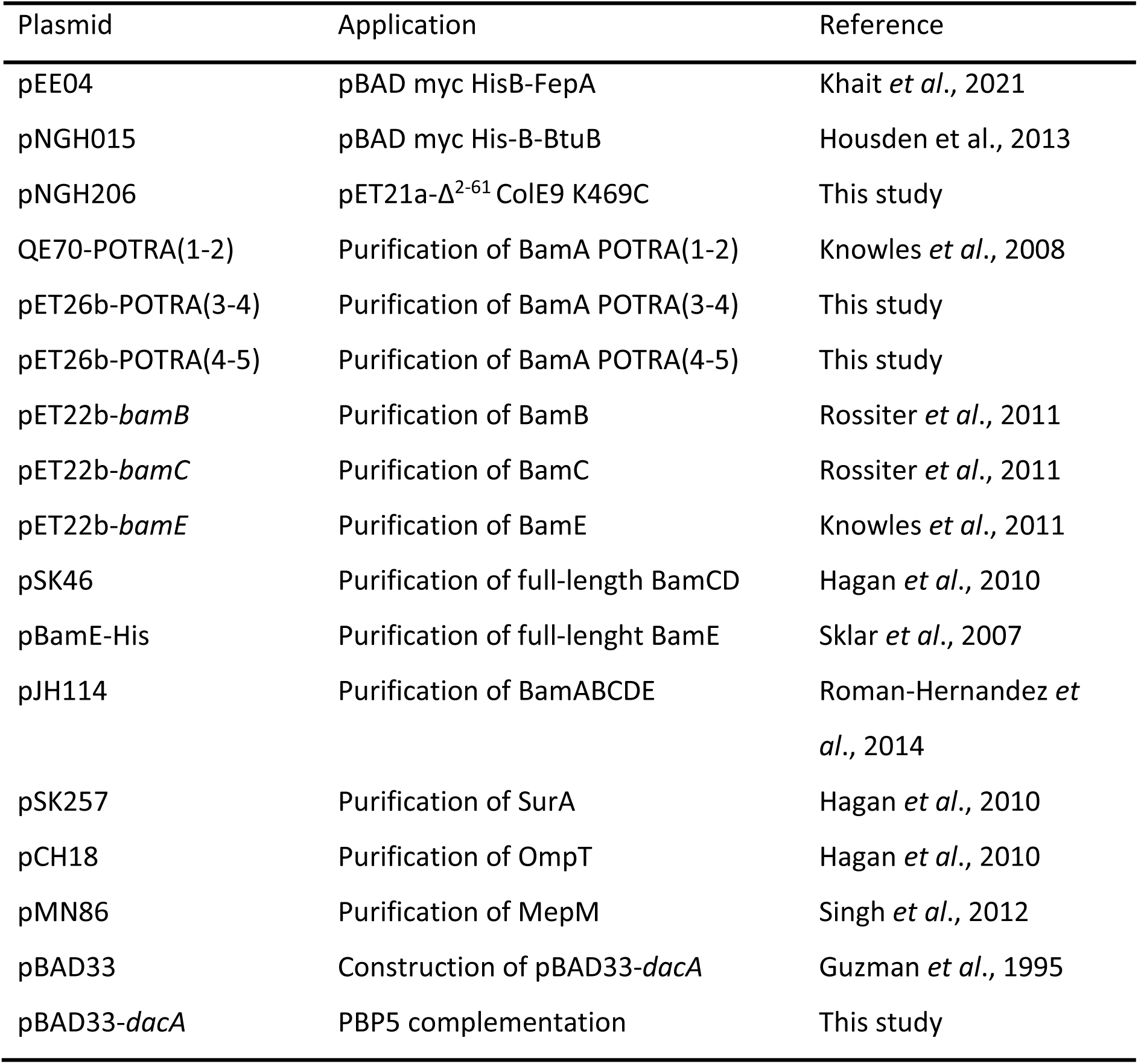
Plasmids used in this study.

**Table S3.**
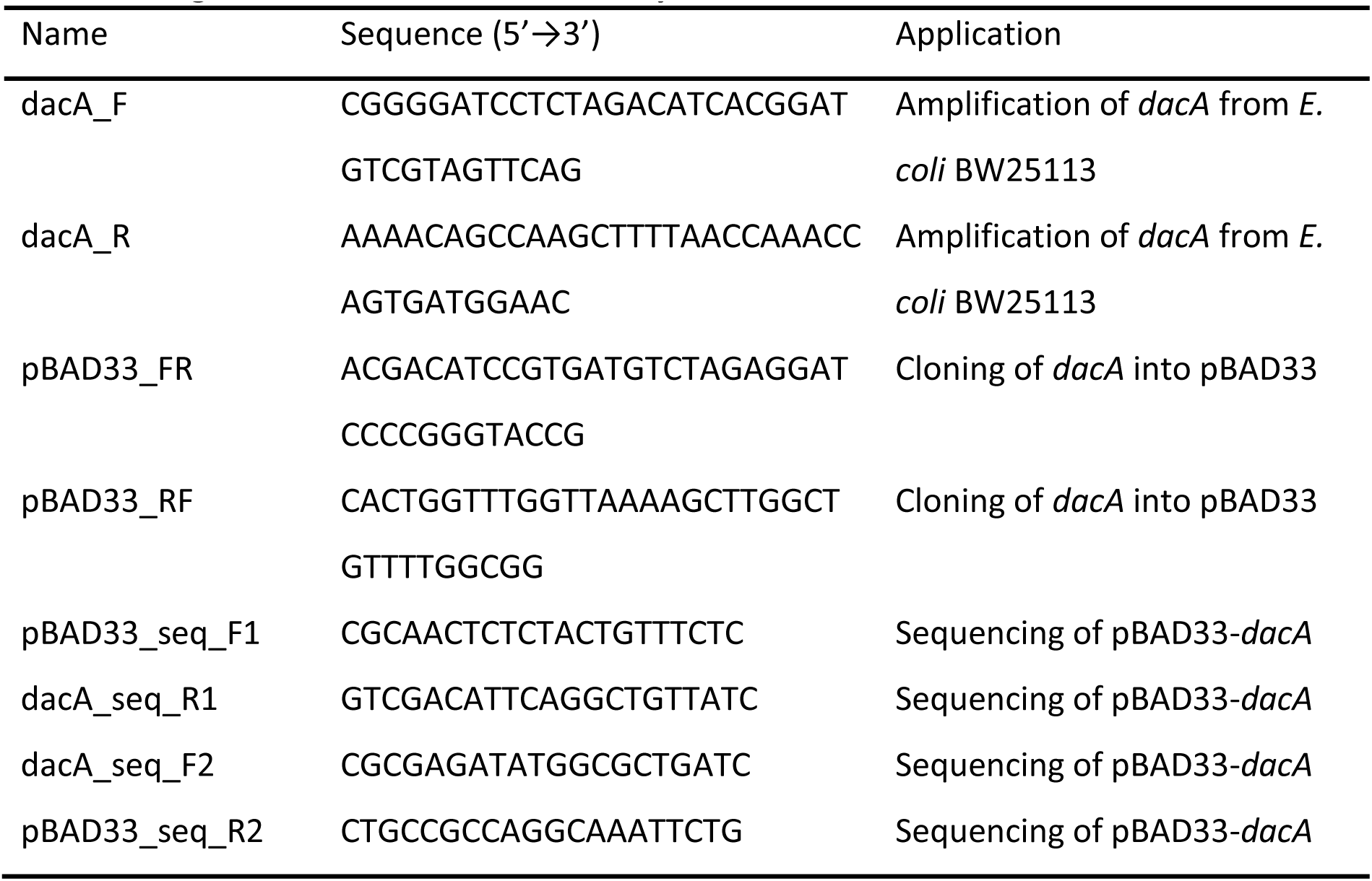
Oligonucleotides used in this study.

**Table S4.**
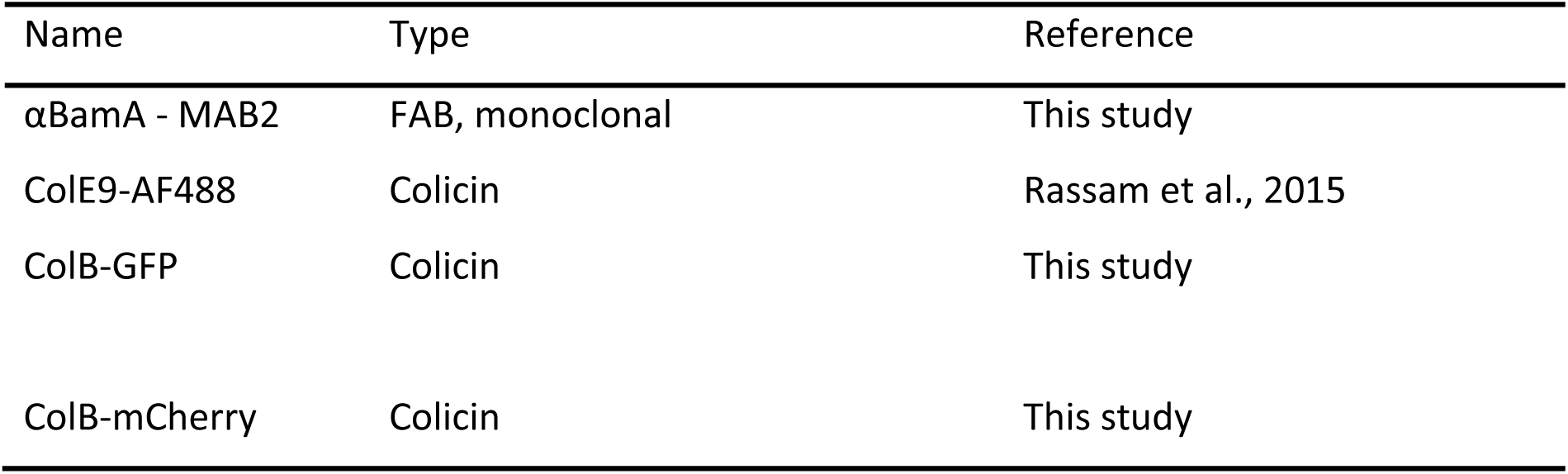
Antibodies and engineered colicins used in this study.

**Extended Data Figure 1.**
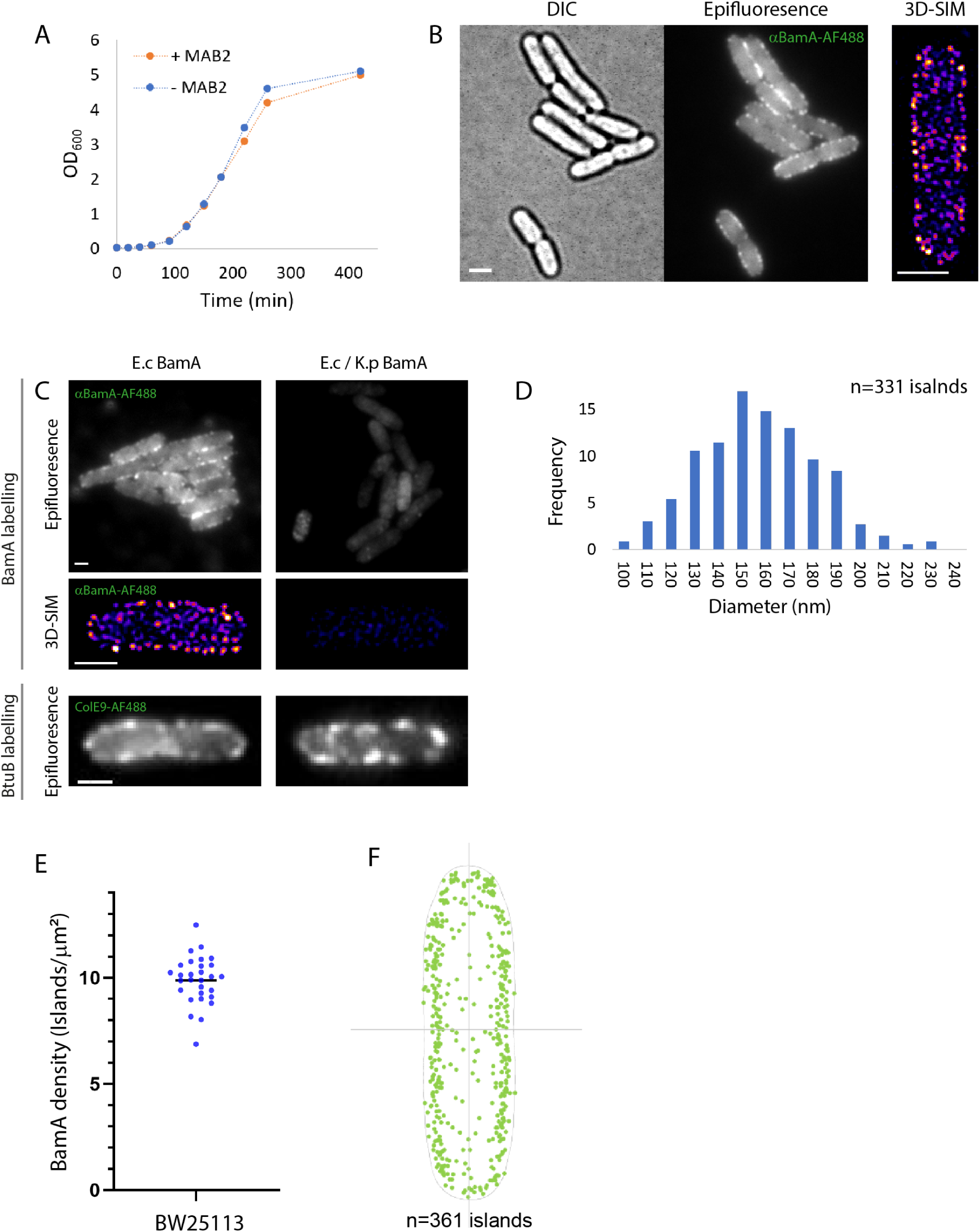
Specific labelling of BamA using high-affinity monoclonal antibodies. **(A)** Growth curves of *E. coli* grown in LB with or without the MAB2 Fabs used for live cell labelling showing the Fabs have no effect on bacterial survival. **(B)** A field of view showing DIC and epifluorescence images of BamA labelled using αBamA^AF488^ Fabs (*left*) and a 3D-SIM image of a single cell (*right*) **(C)** BamA labelling by MAB2 is specific for *E. coli* BamA. BamA and BtuB, used as a control, labelling of the BW25113 strain (*E. coli* sequence) and a strain expressing a modified BamA barrel domain (*E. coli*/*K. pneumoniae* sequence). Shown are comparative images of cells labelled with αBamA^AF488^ or ColE9^AF488^. **(D)** Size distribution of BamA-containing islands demonstrating that the average island diameter is ∼ 150 nm. **(E)** The density of BamA-containing islands on the surface of exponentially growing *E. coli* as measured by 3D-SIM (n=30 cells). **(F)** Integrated localization data from multiple cells showing BamA islands are distributed through the *E. coli* OM. Scale bars, 1 μm.

**Extended Data Figure 2.**
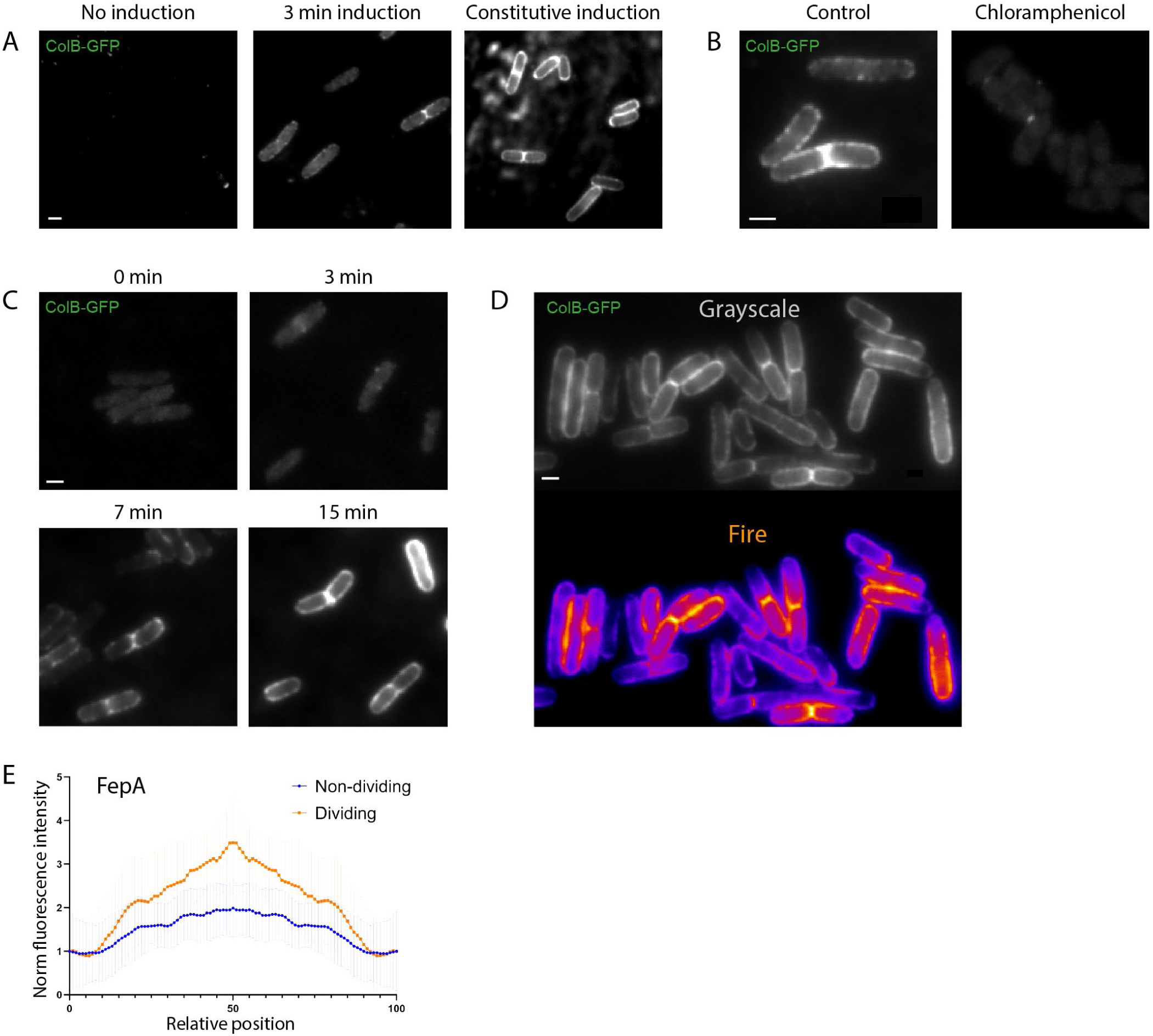
FepA induction system reveals the patterns of OMP biogenesis. **(A)** Epifluorescence images of FepA in an inducible expression strain (GM07). Shown are images of cells after different induction regimes. FepA expression was induced with 0.4% arabinose and the cells stained with ColB-GFP. **(B)** Images of FepA labelling after 7 min induction with or without chloramphenicol pre-treatment demonstrating that labelling is dependent on new synthesis. **(C)** Timeline of FepA biogenesis in M9 media. Samples were taken and labelled with ColB-GFP at different time points after FepA induction. **(D)** A field of view showing grayscale and a corresponding fire heatmap of FepA labelling after 7 min induction. **(E)** The fluorescence distribution (± SD) of FepA labelling in dividing *vs*. non-dividing cells after 7 min induction (n=25 cells each). Scale bars, 1 μm.

**Extended Data Figure 3.**
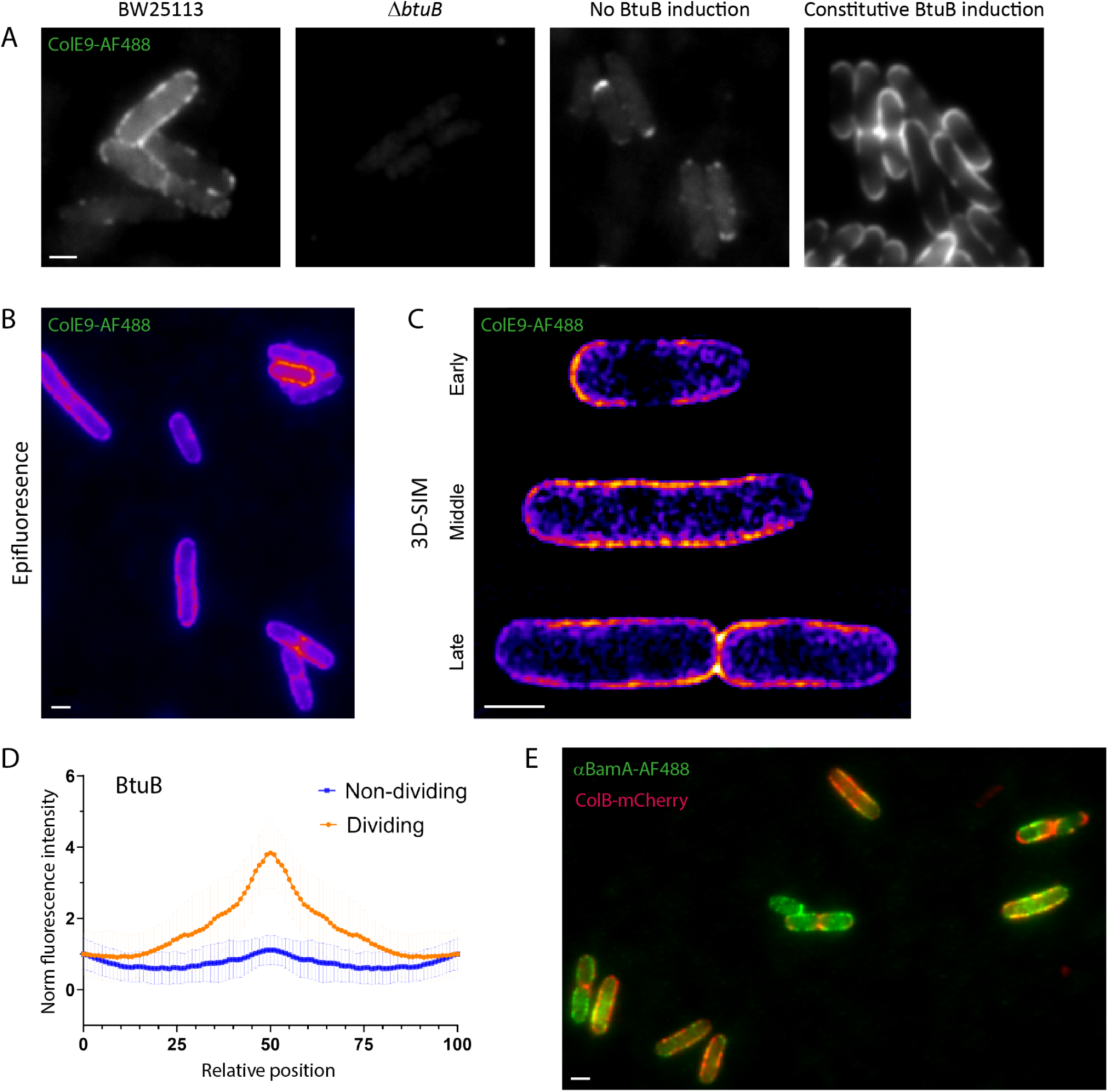
The biogenesis pattern of BtuB matches that of FepA. **(A)** Epifluorescence images of BtuB stained with ColE9^AF488^ under different induction regimes and showing labelling is specific for BtuB. Expression was induced with 0.4% arabinose. **(B)** Representative image of BtuB biogenesis after 5 min induction and staining with ColE9^AF488^. Shown is a heatmap of a field of view. **(C)** 3D-SIM projections of BtuB biogenesis after 5 min induction. The images represent different cell cycle stages. **(D)** The fluorescence distribution (± SD) of BtuB labelling in dividing vs non-dividing cells after 5 min induction (n=30). **(E)** Co-labelling of BamA and FepA with αBamA^AF488^ and ColB-GFP, respectively, after 7 min induction of FepA biogenesis. Shown is an overlay of the BamA (*green*) and FepA (*red*) channels. Scale bars, 1 μm.

**Extended Data Figure 4.**
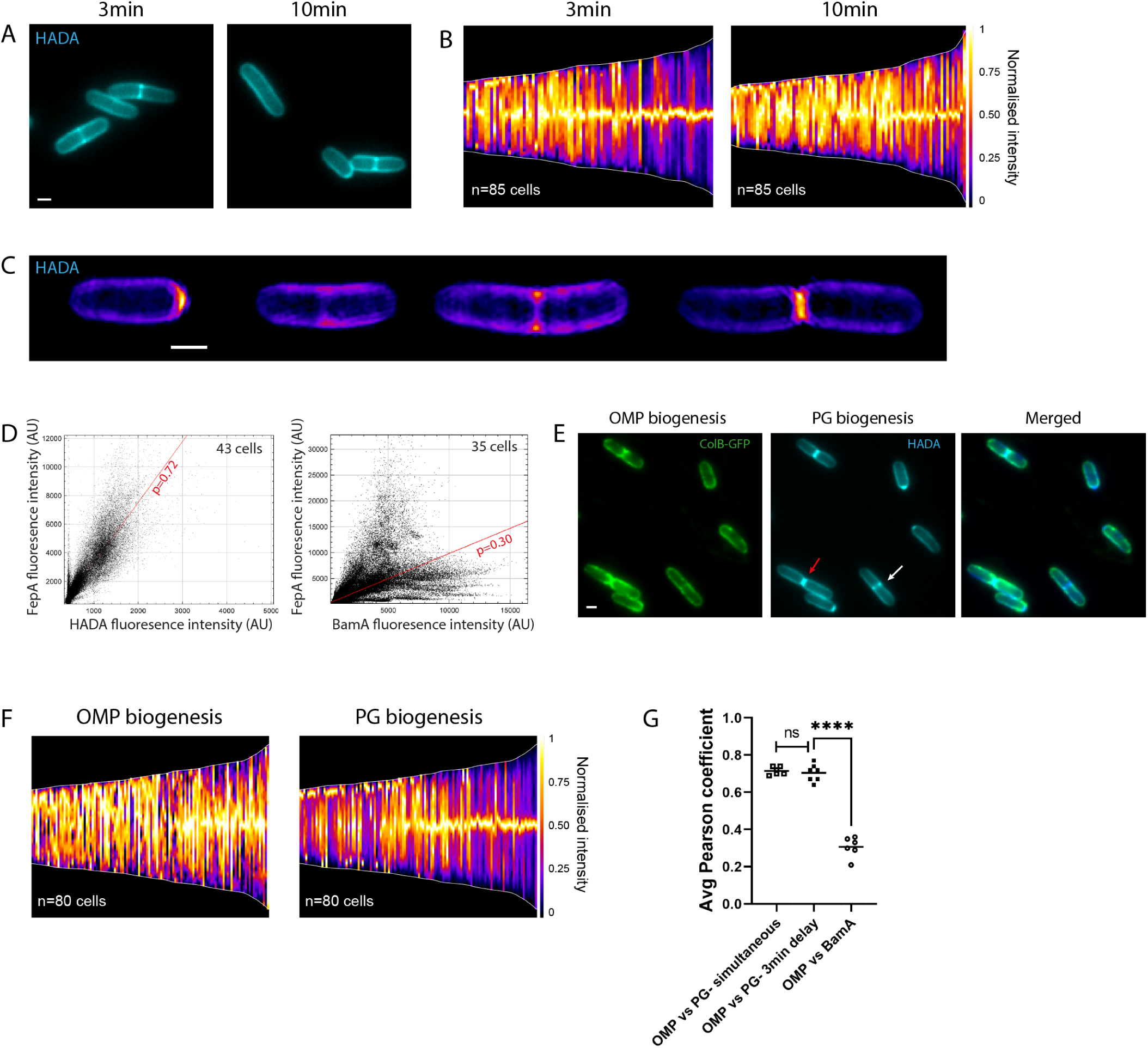
The spatiotemporal organization of cell wall biogenesis mirrors OMP biogenesis. **(A)** Images of *E. coli* cells (BW25113) after incubation with the PG stain HADA for 3 or 10 min. **(B)** Demographs of the normalised fluorescence intensity across the long axis of multiple cells after 3 or 10 min incubation with HADA. Cells are aligned so the more intense pole appears at the top. The charts highlight the cell cycle dependent patterns of PG biogenesis. **(C)** 3D-SIM images of single *E. coli* cells after incubation with HADA for 10 min. *Left to right*, cells representing different stages of the cell cycle. **(D)** The fluorescence intensity of FepA vs HADA (*left*) or BamA (*right*) following simultaneous co-labelling. The representative pixel by pixel cytofluorograms of single images show that OMP biogenesis correlates better with PG biogenesis than BAM localization. **(E)** Co-labelling of PG and OMP biogenesis incorporating a time delay. HADA was added 3 min after FepA induction (stained with ColB-GFP). The total induction time was 10 min. The white/red arrows indicate cells from group1/2 respectively. **(F)** Demographs comparing the normalised fluorescence distribution of FepA and PG biogenesis in multiple cells. Cells were treated as in E **(G)** Histograms showing correlation between FepA biogenesis and either PG biogenesis or BamA distribution. Shown are average Pearson coefficients (+ SD) from 7 images each. Scale bars, 1 μm.

**Extended Data Figure 5.**
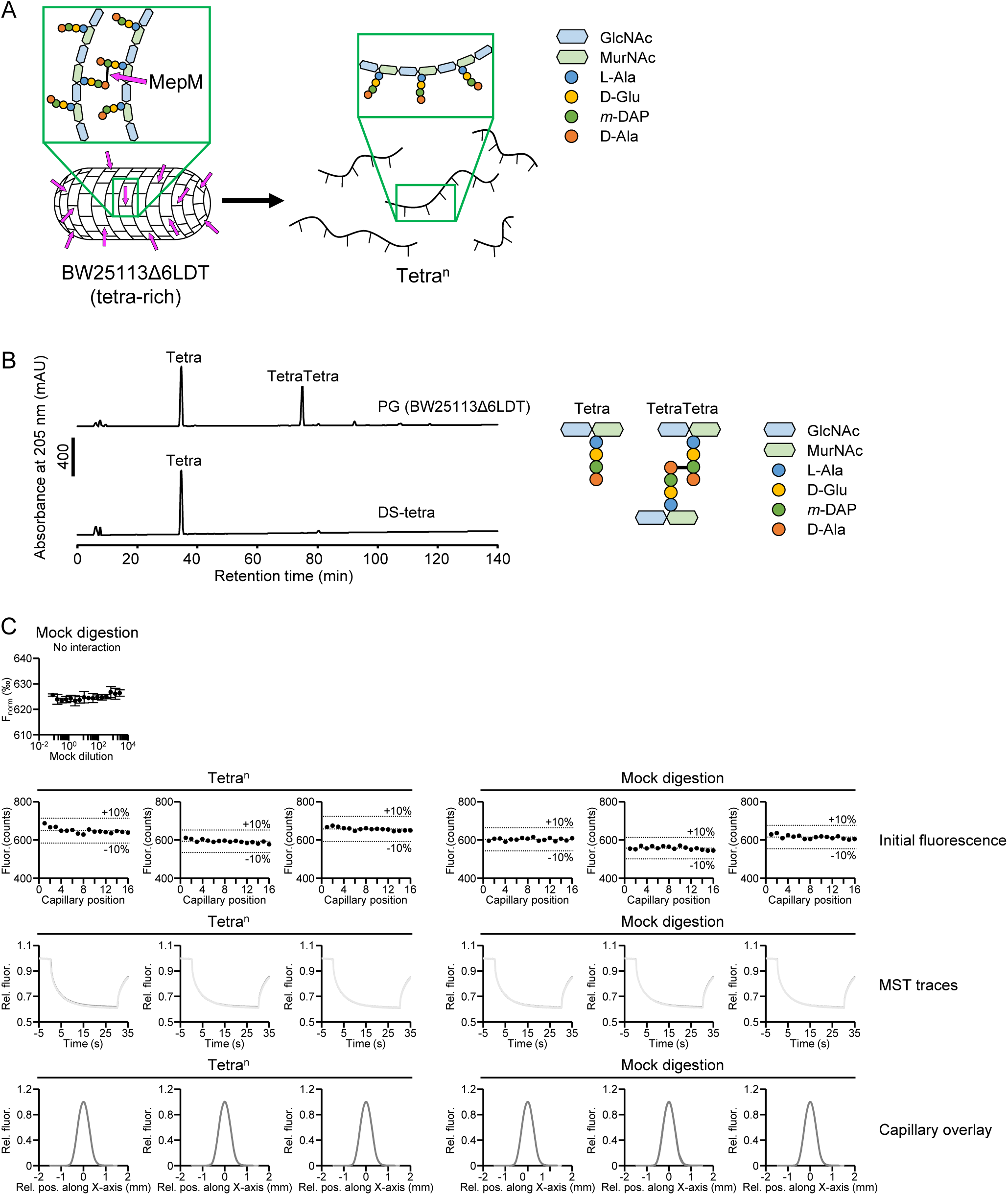
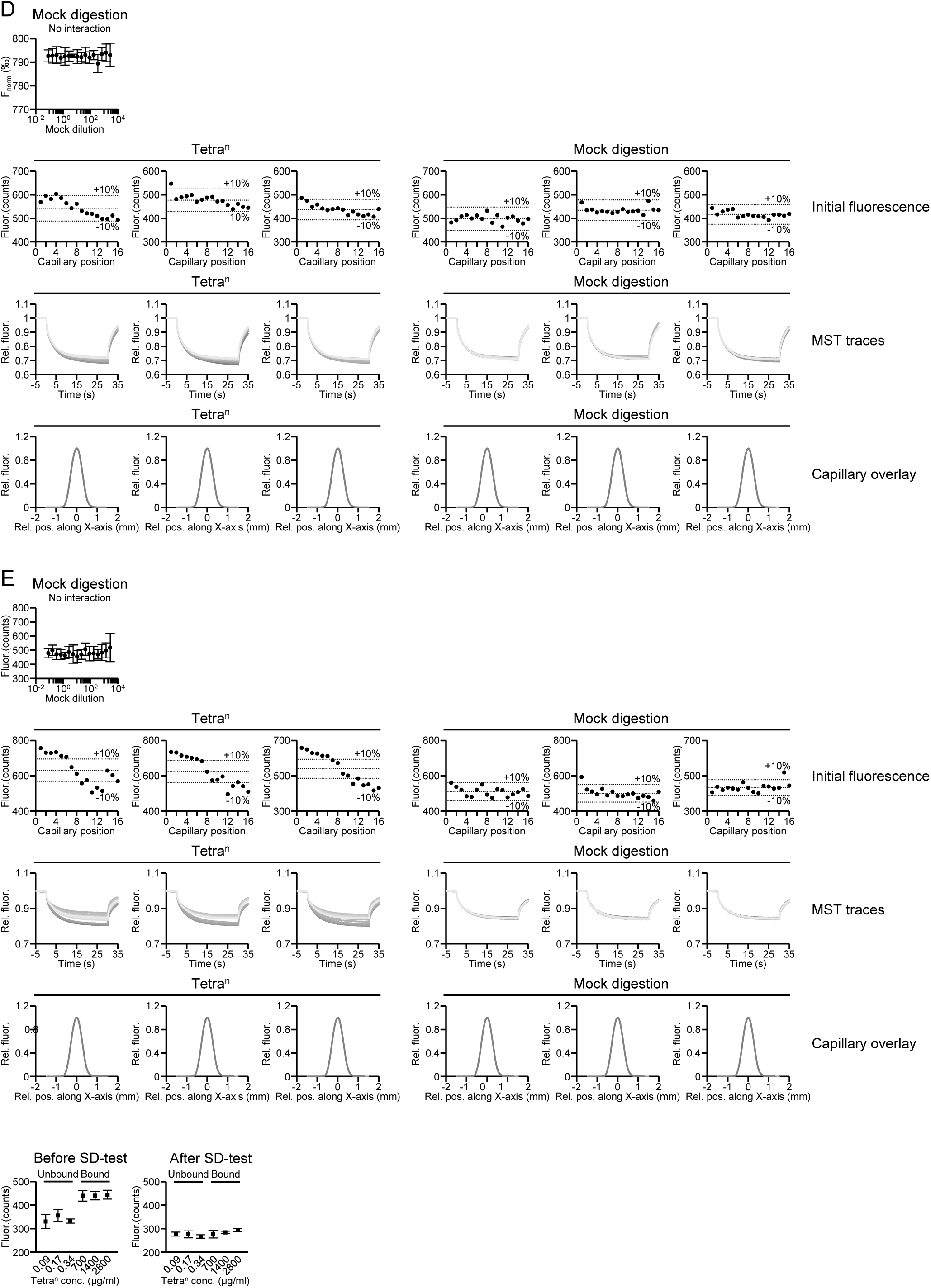
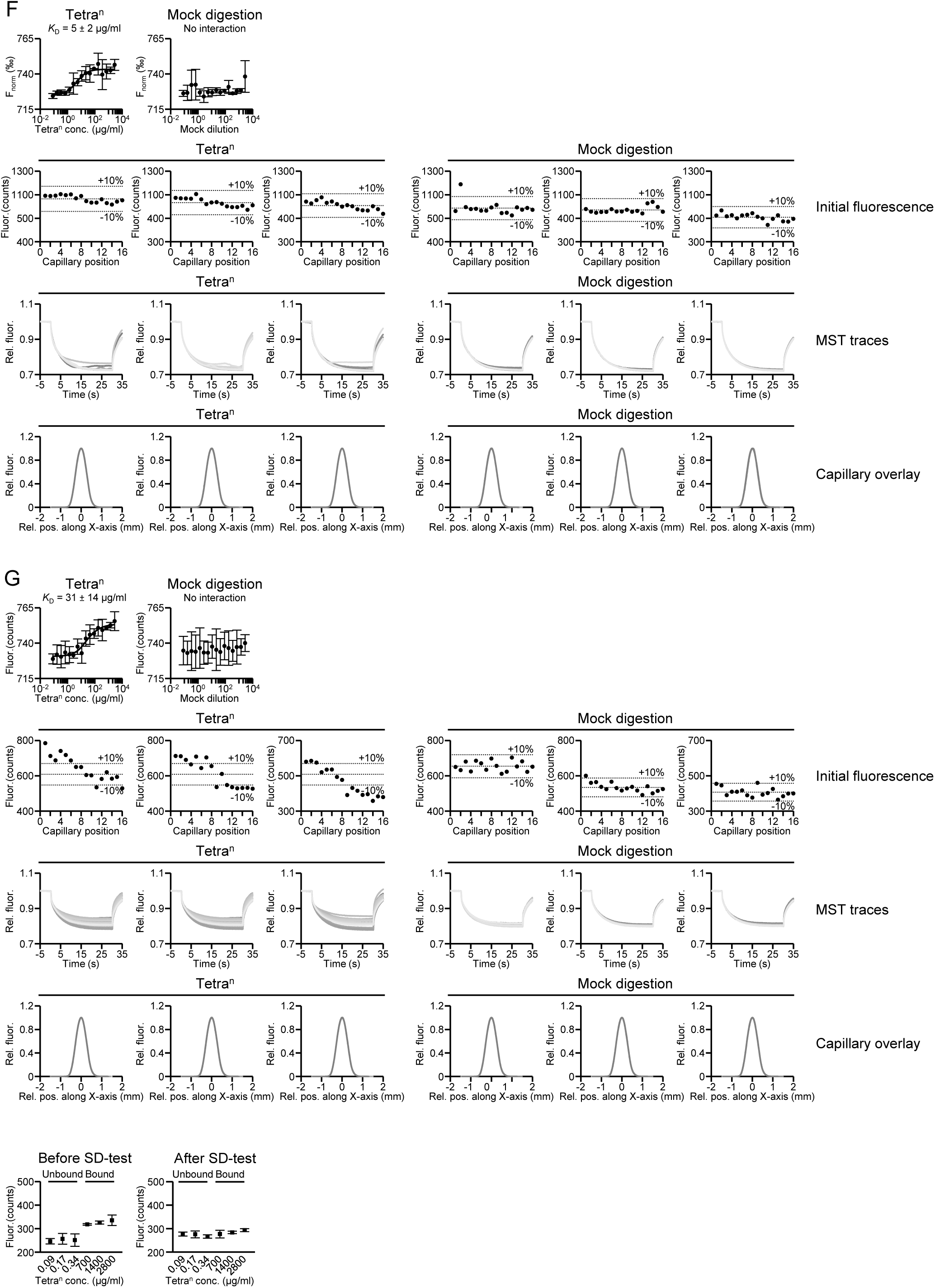
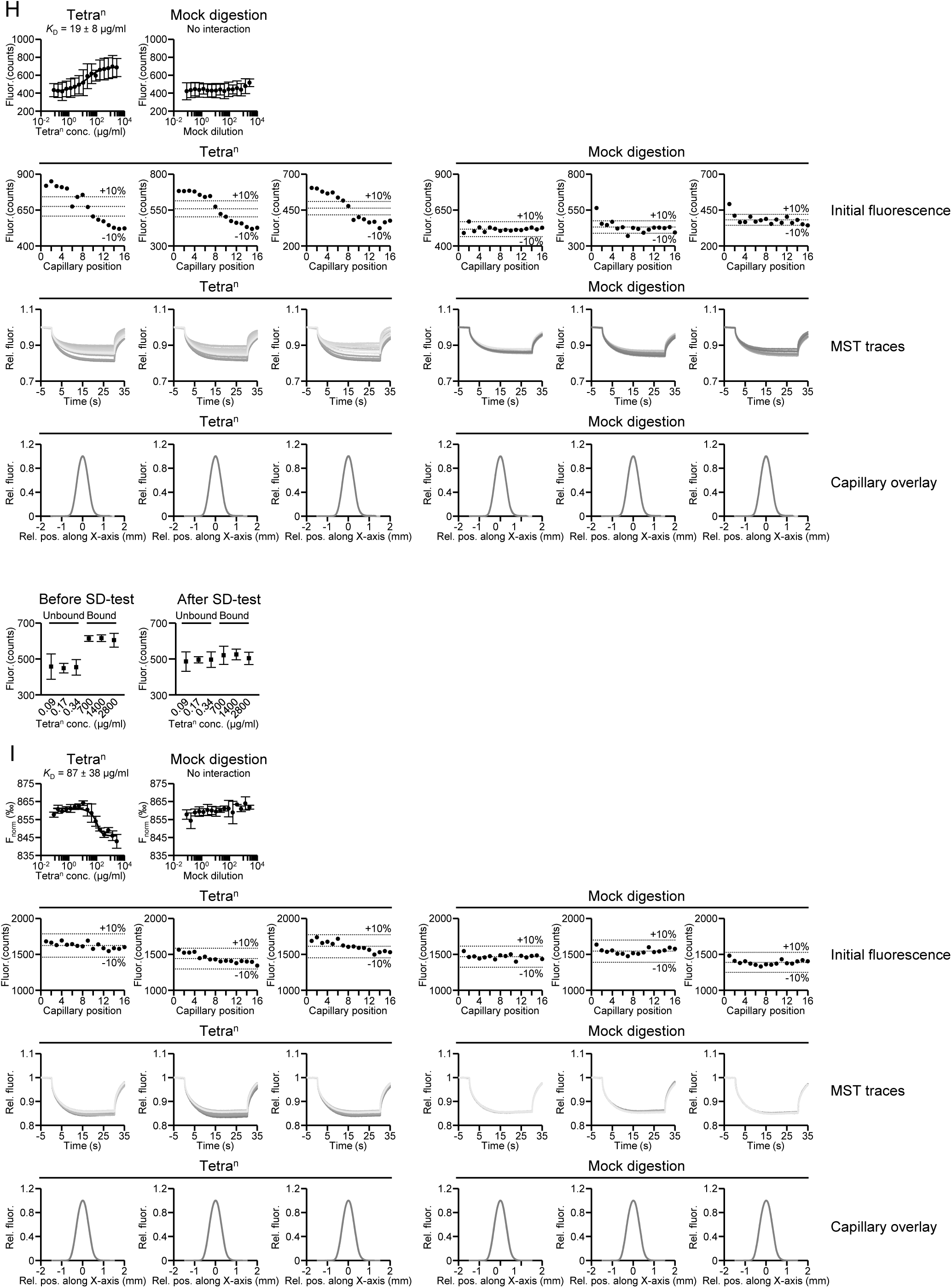
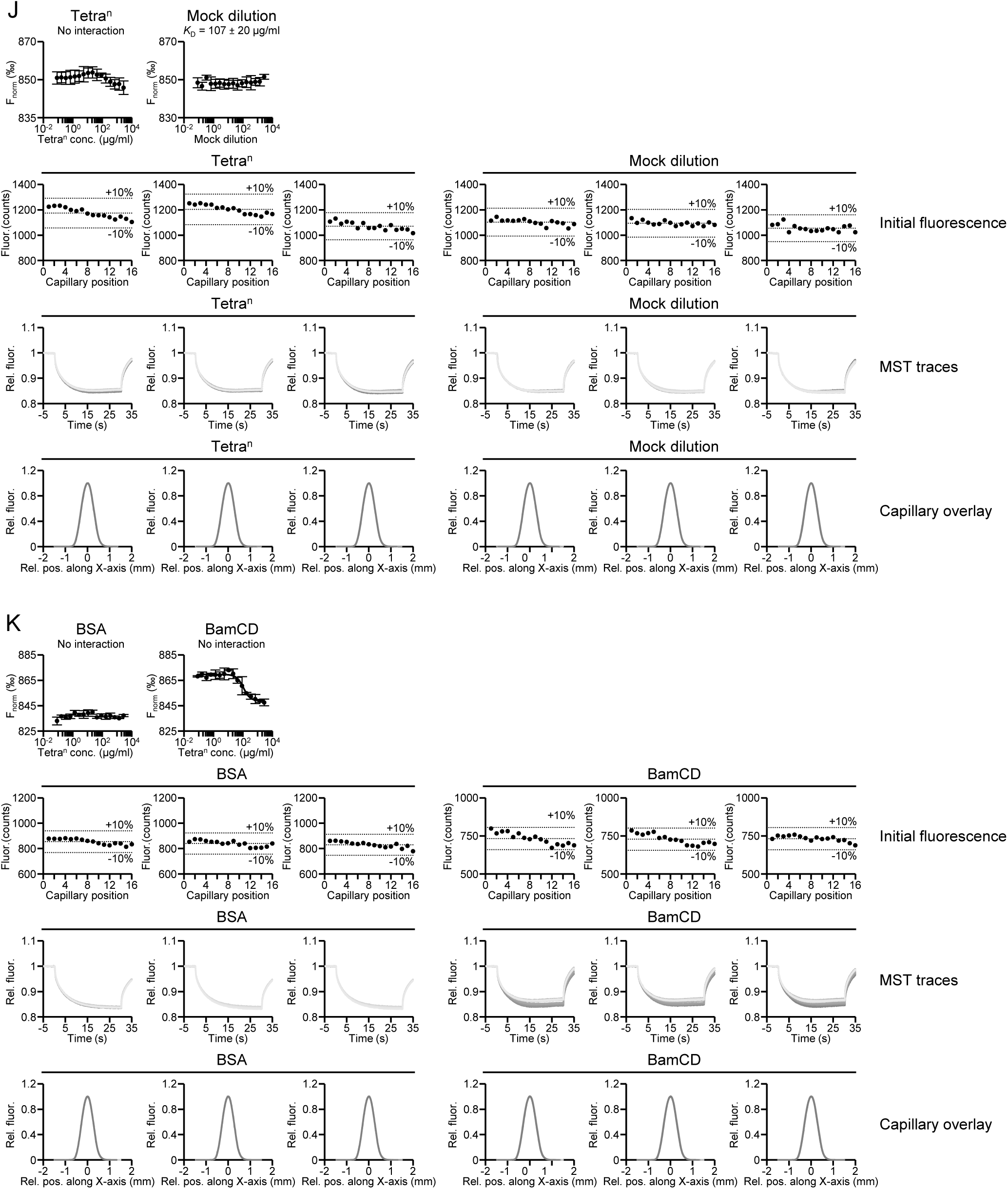
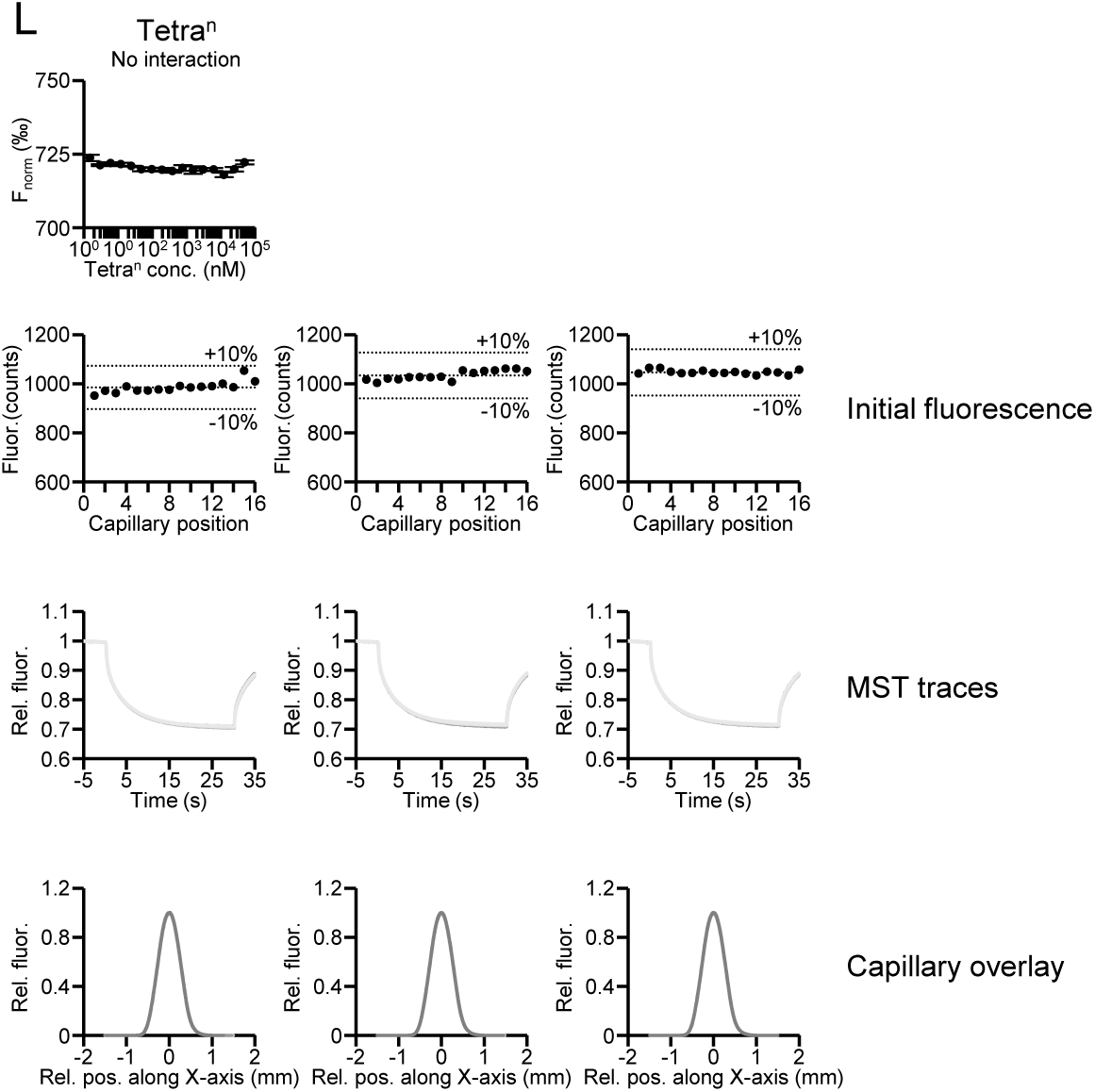
MST experiments for Bam proteins in the presence of Tetra^n^. **(A)** Generation of Tetra^n^ by enzymatic digestion of sacculi from BW25113Δ6LDT. The DD-endopeptidase MepM cleaves the DD-cross-links, generating soluble Tetra^n^ chains of variable length. **(B)** HPLC analysis of sacculi from BW25113Δ6LDT (*top*) and of Tetra^n^ (*bottom*), after digestion with the muramidase cellosyl to produce the disaccharide peptide subunits (muropeptides). Schematic representations of the chemical structures of muropeptides are shown on the right side. **(C)** MST experiment for BamA P1,2 and Tetra^n^. Fig. 2F shows that BamA P1,2 does not interact with Tetra^n^. Here, the top graph shows the lack of MST response in the control sample (mock PG digest without Tetra^n^). The bottom panels show initial fluorescence (average fluorescence ± 10% variation is indicated as dotted lines for each replicate), MST traces and capillary overlay from three independent experiments with Tetra^n^ or the mock PG digest. (**D**) MST experiment as in C but with BamA P3,4. BamA P3,4 interacts with Tetra^n^ (Fig. 2F). (**E**) MST experiment as in C but with BamA P4,5. BamA P4,5 interacts with Tetra^n^ (Fig. 2F). Further MST experiments were performed and initial fluorescence data, MST traces and capillary overlay from three independent experiments in the presence or absence of Tetra^n^ or mock PG digests for BamB **(F)**, BamC **(G)**, BamE **(H)**, BamCD **(I)** and BamCDE **(J)**. Experiments in which the apparent *K*_D_ was determined from ligand-dependent changes in initial fluorescence in the presence of Tetra^n^, i.e. BamA P4,5 **(E)**, BamC **(G)** and BamE **(H)** were validated by SDS-denaturation (SD) tests whereby proteins were or were not denatured prior to the experiment. Values are mean ± SD of three independent experiments. **(K)** Control experiments performed with fluorescent-labelled BSA or BamCD in the presence of Tetra^n^ or mock PG digest, showing that proteins do not generally bind to Tetra^n^ in MST experiments. **(L)** Control experiments performed with free fluorescent Red-NHS dye and no protein in the presence of Tetra^n^, showing that the fluorescent dye does not bind to Tetra^n^.

**Extended Data Figure 6.**
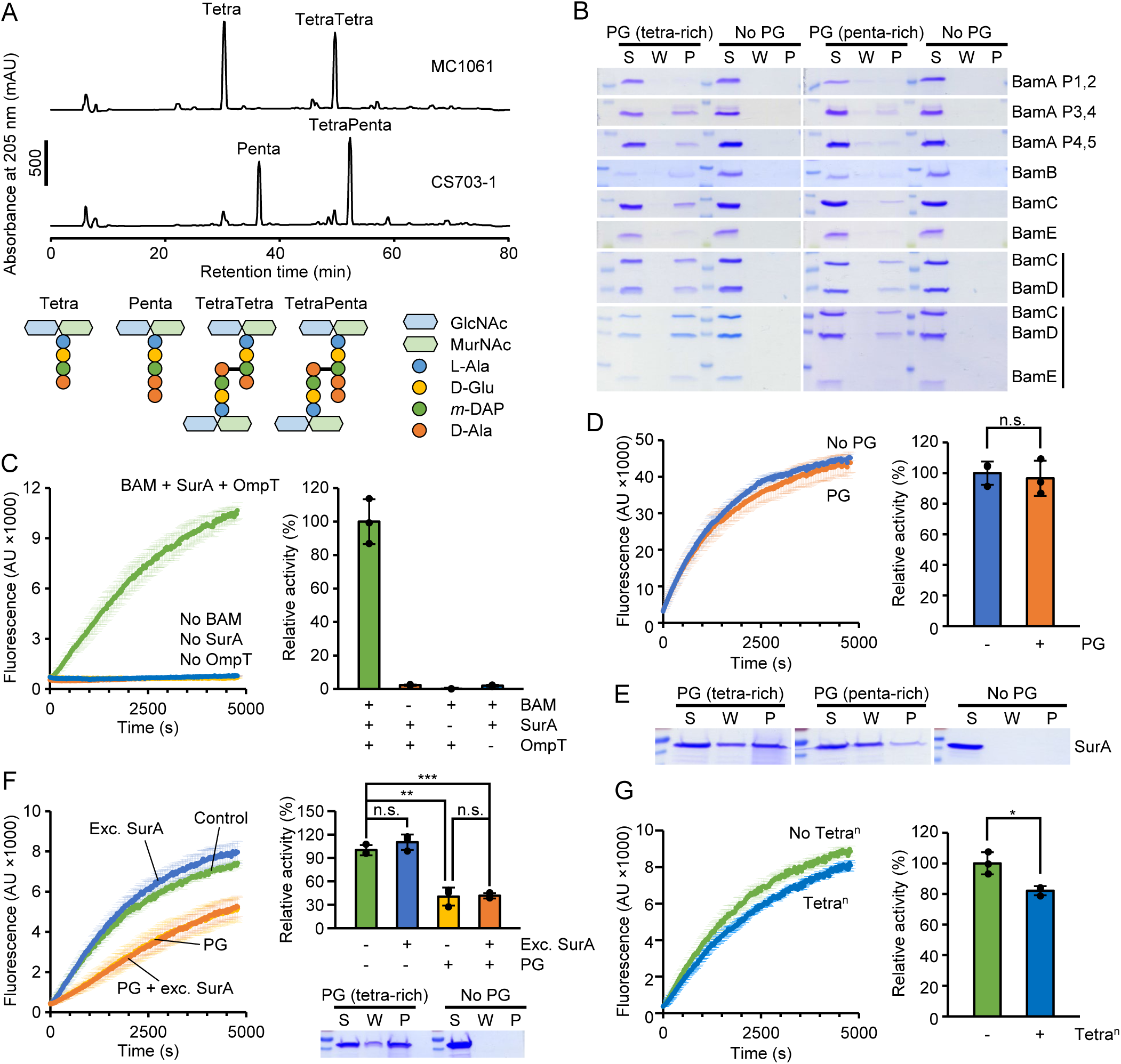
Controls for PG pull-down experiments and BAM activity assays performed with tetrapeptide-rich and pentapeptide-rich PG. **(A)** HPLC analysis of tetrapeptide-rich PG from MC1061 and pentapeptide-rich PG from CS703-1 after digestion with cellosyl to generate the muropeptides. Schematic representation of the main muropeptides are shown. **(B)** PG pull-down assays for Bam proteins performed in the presence of tetrapeptide-rich PG (MC1061) or pentapeptide-rich PG (CS703-1), analysed by SDS-PAGE and Coomassie Blue staining. *S*, supernatant fraction; *W*, wash fraction; *P*, pellet fraction. **(C)** BAM-mediated OmpT assembly *in vitro*. Control experiments without BAM (empty liposomes), OmpT or SurA are included. Values are mean ± SD of three independent experiments. **(D)** Effect of tetrapeptide-rich PG on OmpT activity. Sacculi did not reduce the cleavage of the fluorogenic peptide when added to reactions containing already folded OmpT. **(E)** Interaction of SurA with tetrapeptide-rich (MC1061) and pentapeptide-rich PG (CS703-1) *in vitro,* analysed by SDS-PAGE and Coomassie Blue staining. *S*, supernatant fraction; *W*, wash fraction; *P*, pellet fraction. **(F)** BAM activity assays performed in the presence of a 15 µM excess of SurA and tetrapeptide-rich PG, showing that SurA binding to PG is not a limiting factor for BAM activity *in vitro*. **(G)** Effect of Tetra^n^ on BAM-mediated OmpT assembly *in vitro*.

**Extended Data Figure 7.**
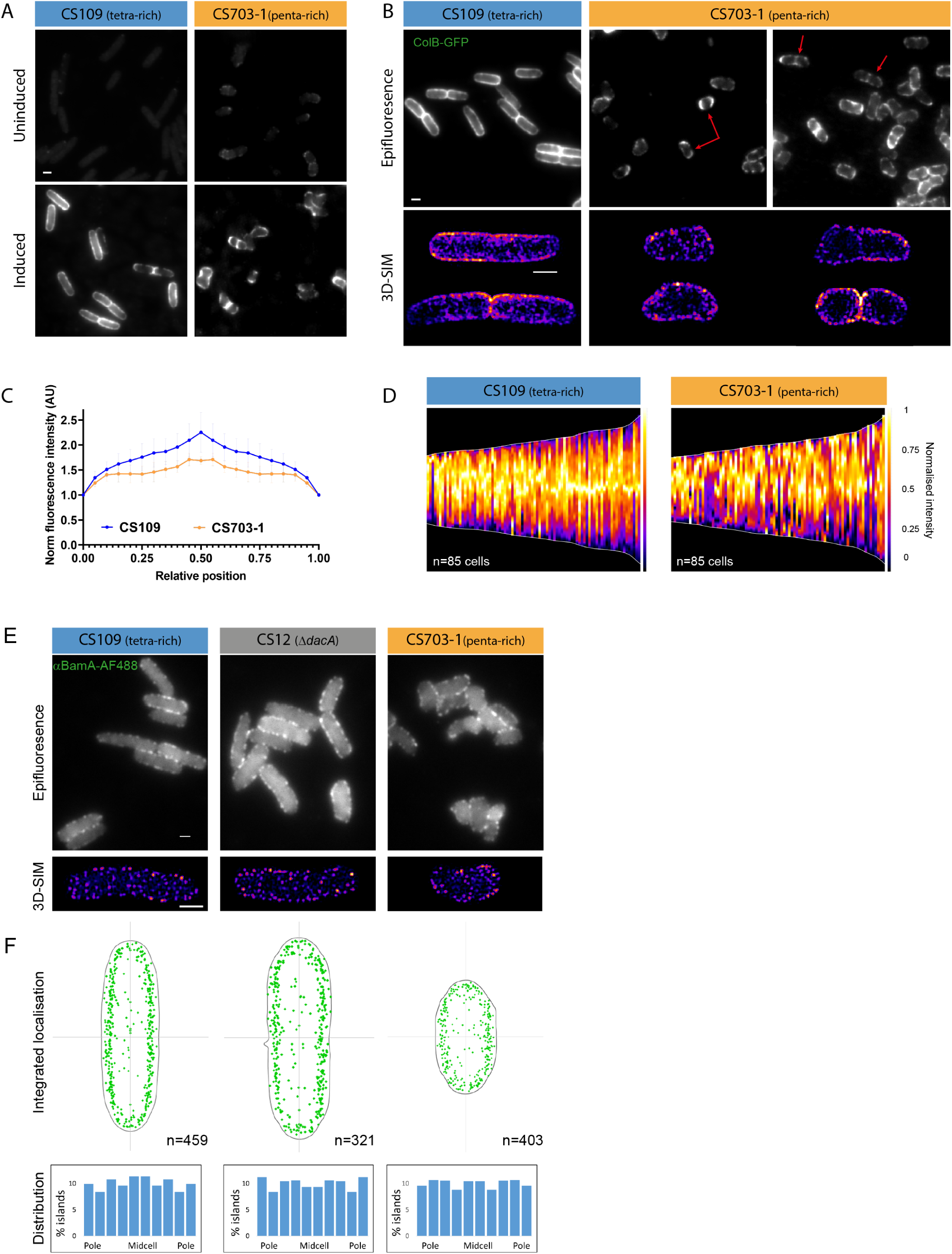
Increased levels of pentapeptide-rich PG diminishes midcell bias of OMP biogenesis. **(A)** Epifluorescence images of FepA, labelled with ColB-GFP in the indicated strains with or without induction of FepA expression. **(B)** Epifluorescence and 3D-SIM Images of newly synthesised FepA in the indicated strains after 7 min induction (0.4% arabinose). Red arrows indicate cells exhibiting atypical biogenesis patterns compared to the CS109 parent strain. **(C)** Distribution of FepA localisation along normalised cell lengths in the strains from B (± SD, 3 biological replicates). **(D)** The normalised fluorescence intensity across the long-axis of multiple cells. Shown are demographs of FepA labelling 7 min after induction of the indicated strains. These charts and the images in B demonstrate the incoherent and less midcell biased OMP biogenesis patterns of the pentapeptide-rich strain. **(E)** Epifluorescence (*top*) and 3D-SIM *(bottom)* images of the indicated strains labelled with αBamA^AF488^ Fabs. The images highlight that BamA organization is similar despite the changes in PG composition. **(F)** Comparison of BamA distribution in the strains from panel C. Shown are integrated localization maps (*top*) and charts showing the distribution of islands along the long axis of the cells (bottom). The charts show that BamA is uniformly distributed in all strains.

**Extended Data Figure 8.**
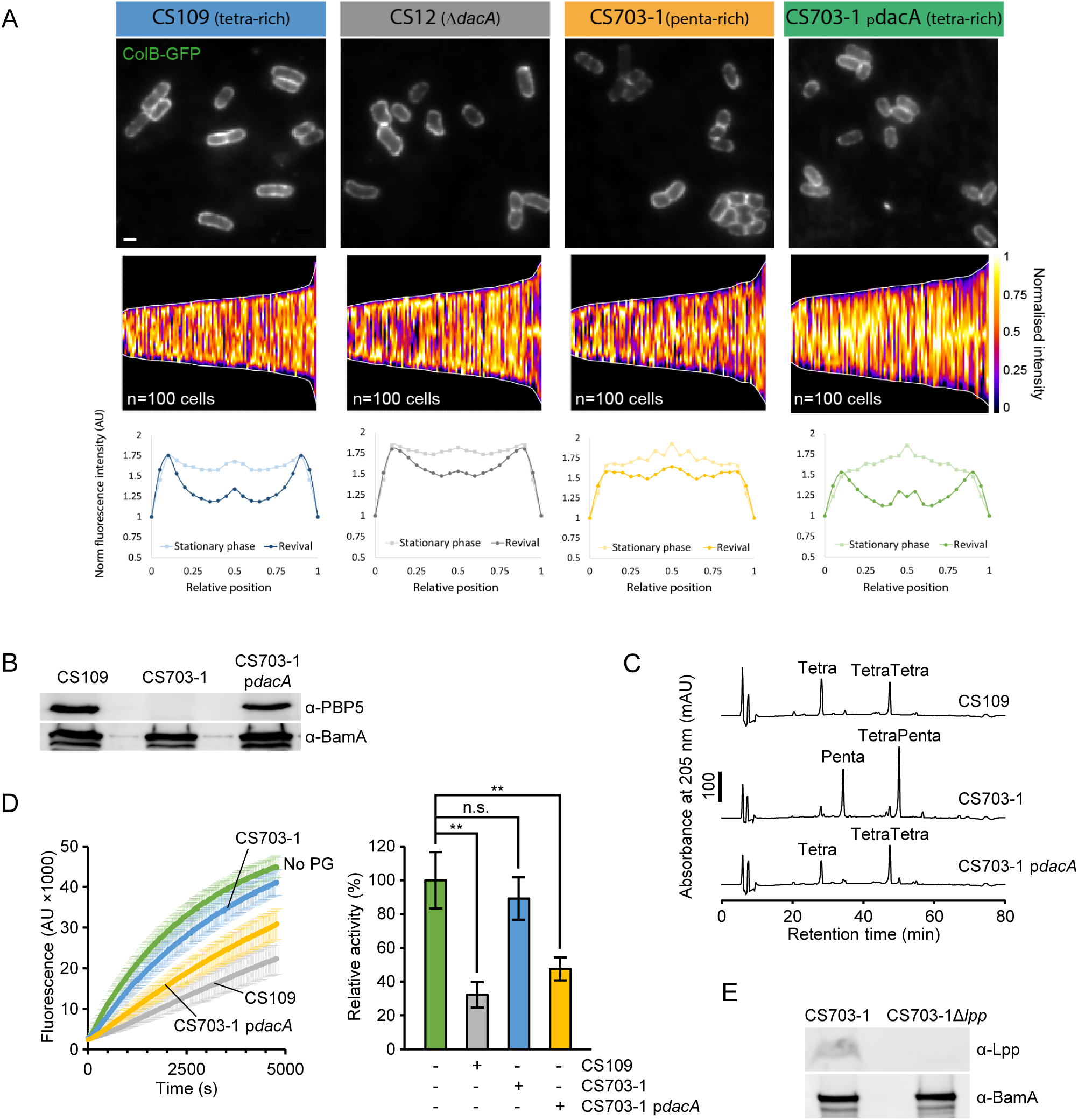
Controls for complementation experiments by ectopic PBP5 production from p*dacA*. **(A)** Native FepA localisation before resuspension in fresh media (stationary phase). Shown are representative images of the indicated strains (*top*), demographs of the normalised fluorescence intensity across the long axis of multiple cells (*middle*) and distribution of native FepA before vs after revival (*bottom*). The charts and images demonstrate that FepA distribution is almost uniform at stationary phase in all strains but binary partitioning is diminished in penta-rich strains. **(B)** Ectopic PBP5 production from p*dacA* in CS703-1 detected by Western Blot and specific antibodies. **(C)** HPLC analysis of cellosyl-digested PG sacculi shows that the CS703-1 cells expressing PBP5 restore a tetrapeptide-rich PG as is present in the CS109 strain. **(D)** Effect of PBP5-remodelled PG isolated from CS703-1 (with or without expression of PBP5) on BAM-mediated OmpT assembly *in vitro*. **(E)** Western Blot analysis confirming the absence of Lpp in CS703-1Δ*lpp*.

**Extended Data Figure 9.**
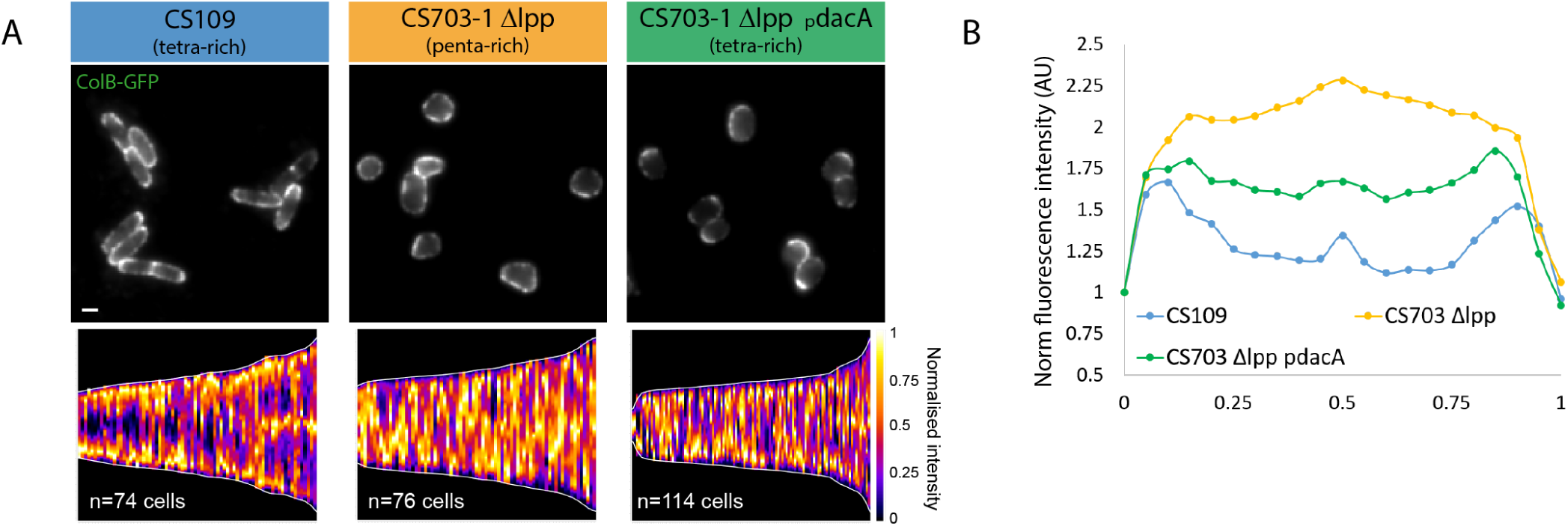
The effect of PG composition on OMP biogenesis is independent of Lpp. **(A)** Polar migration of pre-existing FepA during revival from stationary phase in the Δ*lpp* background. Shown are representative images (*top*) and demographs of the normalised fluorescence intensity across multiple cells (*bottom*) 45 min after resuspension in fresh M9 media. **(B)** Distribution of FepA localisation along normalised cell lengths in the strains from A. The charts demonstrate that *dacA* expression restores FepA polar migration in the pentapeptide-rich strain in Δ*lpp* background

## Videos Legends

**Video 1** - 3D-SIM image of BamA labelling using monoclonal Fabs (heatmap) and the binary map of the islands.

**Video 2** - 3D-SIM image of FepA labelling after 5 minutes induction (heatmap) in dividing and non-dividing cells.

**Video 3** - 3D-SIM image of PG labelling after 7 minutes incubation with HADA in dividing and non-dividing cells.

## References

1. Braun, V. & Bosch, V. Repetitive sequences in the murein-lipoprotein of the cell wall of Escherichia coli. Proc. Natl. Acad. Sci. U. S. A. 69, 970–974 (1972).

2. Silhavy, T. J., Kahne, D. & Walker, S. The Bacterial Cell Envelope. Cold Spring Harb Perspect Biol 2, 1–16 (2010).

3. Rojas, E. R. et al. The outer membrane is an essential load-bearing element in Gram- negative bacteria. Nature 559, 617–621 (2018).

4. Szczepaniak, J. et al. The lipoprotein Pal stabilises the bacterial outer membrane during constriction by a mobilisation-and-capture mechanism. Nat. Commun. 11, 112–114 (2020).

5. Godessart, P. et al. β-Barrels covalently link peptidoglycan and the outer membrane in the α-proteobacterium Brucella abortus. Nat. Microbiol. 6, 27–33 (2021).

6. Voulhoux, R., Bos, M. P., Geurtsen, J., Mols, M. & Tommassen, J. Role of a highly conserved bacterial protein in outer membrane protein assembly. Science 299, 262–265 (2003).

7. Wu, T. et al. Identification of a multicomponent complex required for outer membrane biogenesis in Escherichia coli. Cell 121, 235–245 (2005).

8. Rassam, P. et al. Supramolecular assemblies underpin turnover of outer membrane proteins in bacteria. Nature 523, 333–336 (2015).

9. Egan, A. J. F., Errington, J. & Vollmer, W. Regulation of peptidoglycan synthesis and remodelling. Nat. Rev. Microbiol. 18, 446–460 (2020).

10. Han, L. et al. Structure of the BAM complex and its implications for biogenesis of outer- membrane proteins. Nat. Struct. Mol. Biol. 23, 192–196 (2016).

11. Gu, Y. et al. Structural basis of outer membrane protein insertion by the BAM complex. Nature 531, 64–69 (2016).

12. Bakelar, J., Buchanan, S. K. & Noinaj, N. The structure of the β-barrel assembly machinery complex. Science 351, 180–186 (2016).

13. Storek, K. M. et al. Monoclonal antibody targeting the β-barrel assembly machine of Escherichia coli is bactericidal. Proc. Natl. Acad. Sci. U. S. A. 115, 3692–3697 (2018).

14. Imai, Y. et al. A new antibiotic selectively kills Gram-negative pathogens. Nature 576, 459– 464 (2019).

15. Luther, A. et al. Chimeric peptidomimetic antibiotics against Gram-negative bacteria. Nature 576, 452–458 (2019).

16. Hart, E. M. et al. A small-molecule inhibitor of BamA impervious to efflux and the outer membrane permeability barrier. Proc. Natl. Acad. Sci. U. S. A. 116, 21748–21757 (2019).

17. Doyle, M. T. & Bernstein, H. D. Bacterial outer membrane proteins assemble via asymmetric interactions with the BamA β-barrel. Nat. Commun. 10, 1–13 (2019).

18. Tomasek, D. et al. Structure of a nascent membrane protein as it folds on the BAM complex. Nature 583, 473–478 (2020).

19. Burgess, N. K., Dao, T. P., Stanley, A. M. & Fleming, K. G. β-Barrel proteins that reside in the Escherichia coli outer membrane in vivo demonstrate varied folding behavior in vitro. J. Biol. Chem. 283, 26748–26758 (2008).

20. Lee, J. et al. Formation of a β-barrel membrane protein is catalyzed by the interior surface of the assembly machine protein BamA. Elife 8, 1–20 (2019).

21. Ursell, T. S., Trepagnier, E. H., Huang, K. C. & Theriot, J. A. Analysis of Surface Protein Expression Reveals the Growth Pattern of the Gram-Negative Outer Membrane. PLOS Comput. Biol. 8, e1002680 (2012).

22. Ryter, A., Shuman, H. & Schwartz, M. Integration of the receptor for bacteriophage lambda in the outer membrane of Escherichia coli: coupling with cell division. J. Bacteriol. 122, 295– 301 (1975).

23. Vos-Scheperkeuter, G. H., Pas, E. & Brakenhoff, G. J. Topography of the insertion of LamB protein into the outer membrane of Escherichia coli wild-type and lac-lamB cells. J. Bacteriol. 159, 440–447 (1984).

24. Smit, J. & Nikaido, H. Outer membrane of gram-negative bacteria. XVIII. Electron microscopic studies on porin insertion sites and growth of cell surface of Salmonella typhimurium. J. Bacteriol. 135, 687–702 (1978).

25. Gunasinghe, S. D. et al. The WD40 Protein BamB Mediates Coupling of BAM Complexes into Assembly Precincts in the Bacterial Outer Membrane. Cell Rep. 23, 2782–2794 (2018).

26. Cohen-Khait, R., Harmalkar, A., Pham, Ph., Webby, M.N., Housden, N.G., Elliston, E., Hopper, J.T.S., Mohammed, S., Robinson, C.V., Gray, J.J. & Kleanthous, C. Colicin-mediated transport of DNA through the iron transporter FepA. MBio In press,.

27. Verwer, R. W. H. & Nanninga, N. Pattern of meso-DL-2,6-diaminopimelic acid incorporation during the division cycle of Escherichia coli. J. Bacteriol. (1980) doi:10.1128/jb.144.1.327-336.1980.

28. Ryter, A., Hirota, Y. & Schwarz, U. Process of cellular division in Escherichia coli growth pattern of E. coli murein. J. Mol. Biol. 78, 185–195 (1973).

29. Kuru, E. et al. Fluorescent D-amino-acids reveal bi-cellular cell wall modifications important for Bdellovibrio bacteriovorus predation. Nat. Microbiol. 2, 1648–1657 (2017).

30. Ursinus, A. et al. Murein (peptidoglycan) binding property of the essential cell division protein FtsN from Escherichia coli. J. Bacteriol. 186, 6728–6737 (2004).

31. Typas, A. et al. Regulation of peptidoglycan synthesis by outer-membrane proteins. Cell 143, 1097–1109 (2010).

32. Singh, S. K., Saisree, L., Amrutha, R. N. & Reddy, M. Three redundant murein endopeptidases catalyse an essential cleavage step in peptidoglycan synthesis of Escherichia coli K12. Mol. Microbiol. 86, 1036–1051 (2012).

33. Li, G. & Peter Howard, S. In Vivo and In Vitro Protein-Peptidoglycan Interactions. Methods Mol. Biol. 1615, 143–149 (2017).

34. Mizuno, T. A Novel Peptidoglycan-Associated Lipoprotein Found in the Cell Envelope of Pseudomonas aeruginosa and Escherichia coli. J. Biochem. 86, 991–1000 (1979).

35. Gray, A. N. et al. Coordination of peptidoglycan synthesis and outer membrane constriction during Escherichia coli cell division. Elife 4, 1–29 (2015).

36. Tomasek, D. & Kahne, D. The assembly of β-barrel outer membrane proteins. Curr. Opin. Microbiol. 60, 16–23 (2021).

37. Meberg, B. M., Sailer, F. C., Nelson, D. E. & Young, K. D. Reconstruction of Escherichia coli mrcA (PBP 1a) mutants lacking multiple combinations of penicillin binding proteins. J. Bacteriol. 183, 6148–6149 (2001).

38. Denome, S. A., Elf, P. K., Henderson, T. A., Nelson, D. E. & Young, K. D. Escherichia coli mutants lacking all possible combinations of eight penicillin binding proteins: Viability, characteristics, and implications for peptidoglycan synthesis. J. Bacteriol. 181, 3981–3993 (1999).

39. Hagan, C. L., Kim, S. & Kahne, D. Reconstitution of outer membrane protein assembly from purified components. Science 328, 890–892 (2010).

40. Roman-Hernandez, G., Peterson, J. H. & Bernstein, H. D. Reconstitution of bacterial autotransporter assembly using purified components. Elife 3, 1–20 (2014).

41. Iadanza, M. G. et al. Lateral opening in the intact β-barrel assembly machinery captured by cryo-EM. Nat. Commun. 7, (2016).

42. Schmidt, A. et al. The quantitative and condition-dependent Escherichia coli proteome. Nat. Biotechnol. 34, 104–110 (2016).

43. Houser, J. R. et al. Controlled Measurement and Comparative Analysis of Cellular Components in E. coli Reveals Broad Regulatory Changes in Response to Glucose Starvation. PLoS Comput. Biol. 11, 1–27 (2015).

44. Potluri, L. et al. Septal and lateral wall localization of PBP5, the major D,D- carboxypeptidase of Escherichia coli, requires substrate recognition and membrane attachment. Mol. Microbiol. 77, 300–323 (2010).

45. Kleanthous, C., Rassam, P. & Baumann, C. G. Protein-protein interactions and the spatiotemporal dynamics of bacterial outer membrane proteins. Curr. Opin. Struct. Biol. 35, 109–115 (2015).

46. Lee, J. et al. Characterization of a stalled complex on the β-barrel assembly machine. Proc. Natl. Acad. Sci. U. S. A. 113, 8717–8722 (2016).

47. Matias, V. R. F., Al-Amoudi, A., Dubochet, J. & Beveridge, T. J. Cryo-transmission electron microscopy of frozen-hydrated sections of Escherichia coli and Pseudomonas aeruginosa. J. Bacteriol. 185, 6112–6118 (2003).

## References

48. Jeong, J. Y. et al. One-step sequence-and ligation-independent cloning as a rapid and versatile cloning method for functional genomics Studies. Appl. Environ. Microbiol. 78, 5440– 5443 (2012).

49. Knowles, T. J., McClelland, D. M., Rajesh, S., Henderson, I. R. & Overduin, M. Secondary structure and 1H, 13C and 15N backbone resonance assignments of BamC, a component of the outer membrane protein assembly machinery in *Escherichia coli*. Biomol. NMR Assign. 3, 203–206 (2009).

50. Knowles, T. J. et al. Fold and function of polypeptide transport-associated domains responsible for delivering unfolded proteins to membranes. Mol. Microbiol. 68, 1216–1227 (2008).

51. Glauner, B. Separation and quantification of muropeptides with high-performance liquid chromatography. Anal. Biochem. 172, 451–464 (1988).

52. Hayashi, K. A rapid determination of sodium dodecyl sulfate with methylene blue. Anal. Biochem. 67, 503–506 (1975).

53. Peters, K., Pazos, M., VanNieuwenhze, M. & Vollmer, W. Optimized Protocol for the Incorporation of FDAA (HADA Labeling) for in situ Labeling of Peptidoglycan. Bio-Protocol 9, 1–12 (2019).

54. Ducret, A., Quardokus, E. M. & Brun, Y. V. MicrobeJ, a tool for high throughput bacterial cell detection and quantitative analysis. Nat. Microbiol. 1, 1–7 (2016).

55. Bolte, S et al., 2011. & Cordelières, F. P. A guided tour into subcellular colocalization analysis in light microscopy. J. Microsc. 224, 213–232 (2006).

56. K. A. Datsenko, B. L. Wanner, One-step inactivation of chromosomal genes in Escherichia coli K-12 using PCR products. Proc. Natl. Acad. Sci. U. S. A. 97, 6640–6645 (2000).

57. K. Heller, B. J. Mann, R. J. Kadner, Cloning and expression of the gene for the vitamin B12 receptor protein in the outer membrane of Escherichia coli. J. Bacteriol. 161, 896–903 (1985).

58. Rossiter, A. E. et al. The essential β-barrel assembly machinery complex components bamD and bamA are required for autotransporter biogenesis. J. Bacteriol. 193, 4250–4253 (2011).

59. Sklar, J. G. et al. Lipoprotein SmpA is a component of the YaeT complex that assembles outer membrane proteins in Escherichia coli. Proc. Natl. Acad. Sci. U. S. A. 21, 2473–2484 (2007).

